# Adaptation of spontaneous activity in the developing visual cortex

**DOI:** 10.1101/2020.07.30.229559

**Authors:** Marina E. Wosniack, Jan H. Kirchner, Ling-Ya Chao, Nawal Zabouri, Christian Lohmann, Julijana Gjorgjieva

## Abstract

Spontaneous activity drives the establishment of appropriate connectivity in different circuits during brain development. In the mouse primary visual cortex, two distinct patterns of spontaneous activity occur before vision onset: local low-synchronicity events originating in the retina, and global high-synchronicity events originating in the cortex. We sought to determine the contribution of these activity patterns to jointly organize network connectivity through different activity-dependent plasticity rules. We found that local events shape cortical input selectivity and topography, while global events have a homeostatic role regulating connection strength. To generate robust selectivity, we predicted that global events should adapt their amplitude to the history of preceding cortical activation, and confirmed by analyzing *in vivo* spontaneous cortical activity. This adaptation led to the sparsification of spontaneous activity on a slower timescale during development, demonstrating the remarkable capacity of the developing sensory cortex to acquire sensitivity to visual inputs after eye-opening.

## Introduction

The impressive ability of the newborn brain to respond to its environment and generate coordinated output without any prior experience suggests that brain networks undergo substantial organization, tuning and coordination even as animals are still in the womb, driven by powerful developmental mechanisms. These broadly belong to two categories: activity-independent mechanisms, involving molecular guidance cues and chemoaffinity gradients which establish the initial coarse connectivity patterns at early developmental stages [1], and activity-dependent plasticity mechanisms which continue the refinement of this initially imprecise connectivity into functional circuits that can execute diverse behaviors in adulthood [2, 3, 4]. Non-random patterns of spontaneous activity drive these refinements, and act as training inputs to the immature circuits before the onset of sensory experience. Many neural circuits in the developing brain generate spontaneous activity, including the retina, hippocampus, cortex and spinal cord (reviewed in [5, 6]). This activity regulates a plethora of developmental processes such as neuronal migration, ion channel maturation and the establishment of precise connectivity [7, 8, 9], and perturbing this activity impairs different aspects of functional organization and axonal refinement [10, 11, 12]. These studies firmly demonstrate that spontaneous activity is necessary and instructive for the emergence of specific and distinct patterns of neuronal connectivity in the developing nervous system.

In the visual system, due to its accessibility, spontaneous activity and its role has been best characterized in the retina where it propagates as waves, and instructs the topographically ordered projection to the thalamus and superior colliculus [7]. This has inspired many computational models of topographic map development based on correlated patterns of activity (reviewed in [13, 14, 15]), including more recent approaches that simulate the entire temporal evolution of map formation from a combination of mechanisms [16, 17]. In the visual, and the sensory cortex more generally, spontaneous activity has been extensively characterized in isolated networks, either in cortical cultures or slices [18, 19, 20, 21]. Here, the spatiotemporal properties of spontaneous activity, including frequency, synchronicity, amplitude and spatial spread, depend on the studied region and developmental age [19, 20, 21, 22]. More recent *in vivo* recordings of endogenous patterns of activity in the developing sensory cortex show that some spontaneous activity properties in the living animal are similar to those described *in vitro* [23, 24, 25]. These studies show that the generation and propagation of spontaneous activity in the intact cortex depend on input from different brain areas. For instance, activity from the sensory periphery substantially contributes to the observed activity patterns in the developing cortex, but there are other independent sources of activity within the cortex itself [24, 26, 27, 25]. Two-photon imaging of spontaneous activity in the *in vivo* mouse primary visual cortex before eye-opening (postnatal days, P8-10) has demonstrated that there are two independently occurring patterns of spontaneous activity with different sources and spatiotemporal characteristics. Peripheral events driven by retinal waves [28, 5] spread in the cortex as local events, engaging a relatively small number of the recorded neurons. In contrast, events intrinsic to the cortex that were unaffected by manipulation of retinal waves spread as global events, synchronously activating a large proportion of the recorded neurons [26].

We know relatively little about the information content of these local and global patterns of spontaneous cortical activity relevant for shaping local and brain-wide neural circuits. Specifically, it is unknown whether spontaneous activity from different sources affects distinct aspects of circuit organization, each providing an independent instructive signal, or if the two patterns cooperate to synergistically guide circuit organization. Therefore, using experimentally characterized properties of spontaneous activity in the visual cortex *in vivo* at P8-10, we developed a biologically plausible, yet analytically tractable, theoretical framework to determine the implications of this activity on normal circuit development with a focus on the topographic refinement of connectivity and the emergence of stable receptive fields.

We investigated the potential of two activity-dependent plasticity rules, the Hebbian covariance rule [29, 30, 31] and the Bienenstock-Cooper-Munro (BCM) rule [32], to generate stable and topographically refined receptive fields driven by L- and H-events. We postulated that peripheral L-events play a key role in topographically organizing receptive fields in the cortex, while H-events regulate connection strength homeostatically, operating in parallel to network refinements by L-events. We considered that H-events are ideally suited for this purpose because they maximally activate many neurons simultaneously, hence lack topographic information that can be used for synaptic refinement. Indeed, under the BCM rule, we found that these H-events dynamically slide the threshold between potentiation and depression, implementing the postulated homeostatic regulation. However, topography is disrupted since small L-events, which carry precise information for topographic connectivity refinements, mostly cause long-term depression in the synaptic weights. In comparison, under the Hebbian rule, we found that connectivity refinements are extremely sensitive to the properties of H-events. Therefore, we proposed that instead of regulating the threshold between potentiation and depression as in the BCM rule, H-events are themselves self-regulating. We implemented this self-regulation in the Hebbian rule through a fast adaptive mechanism that modulates H-event amplitude relative to ongoing activity in the network, effectively adjusting the homeostatic influence of H-events on connection strength. Re-examining the same spontaneous activity that inspired the model [26], we found a robust signature of adaptation in the amplitude of the cortically generated events. In our model, this adaptation led to the sparsification of cortical spontaneous activity over a prolonged timescale of development as observed experimentally in the visual and somatosensory cortices [33, 34], and previously unaccounted by other computational models. Therefore, our work proposes that global, cortically generated activity in the form of H-events rapidly adapts to ongoing network activity, supporting topographic organization of connectivity and maintaining synaptic strengths in an operating regime. The same adaptive mechanism eventually leads to the break up of large-scale spontaneous events, which acts to retain sensitivity of the cortex to incoming peripheral inputs as it transitions towards being visually-driven.

## Results

### A network model for connectivity refinements driven by spontaneous activity

Spontaneous activity in the mouse primary visual cortex before eye-opening exhibits two independently occurring patterns of spontaneous activity (Fig. 1A): (1) local low-synchronicity events involving 20-80% of the recorded neurons originating from the retina and propagated through the thalamus, called L-events and (2) global high-synchronicity events intrinsic to the cortex that activate nearly all cortical neurons (80-100%), called H-events [26]. This classification of spontaneous events into L- and H-events was recently validated both at the single-cell and population levels [35]. We first re-analyzed the extracted events from the original publication [26] and confirmed differences in specific features of these spontaneous events, as well as characterized novel aspects (Fig. 1B). In particular, L-events have lower amplitudes with smaller variance and inter-event intervals (IEI, the inverse of firing frequency) that follow an exponential-like distribution. H-events have a broader amplitude distribution with higher values and IEIs that follow a long-tail distribution. We found that L- and H-events have similar durations.

**Figure 1.**
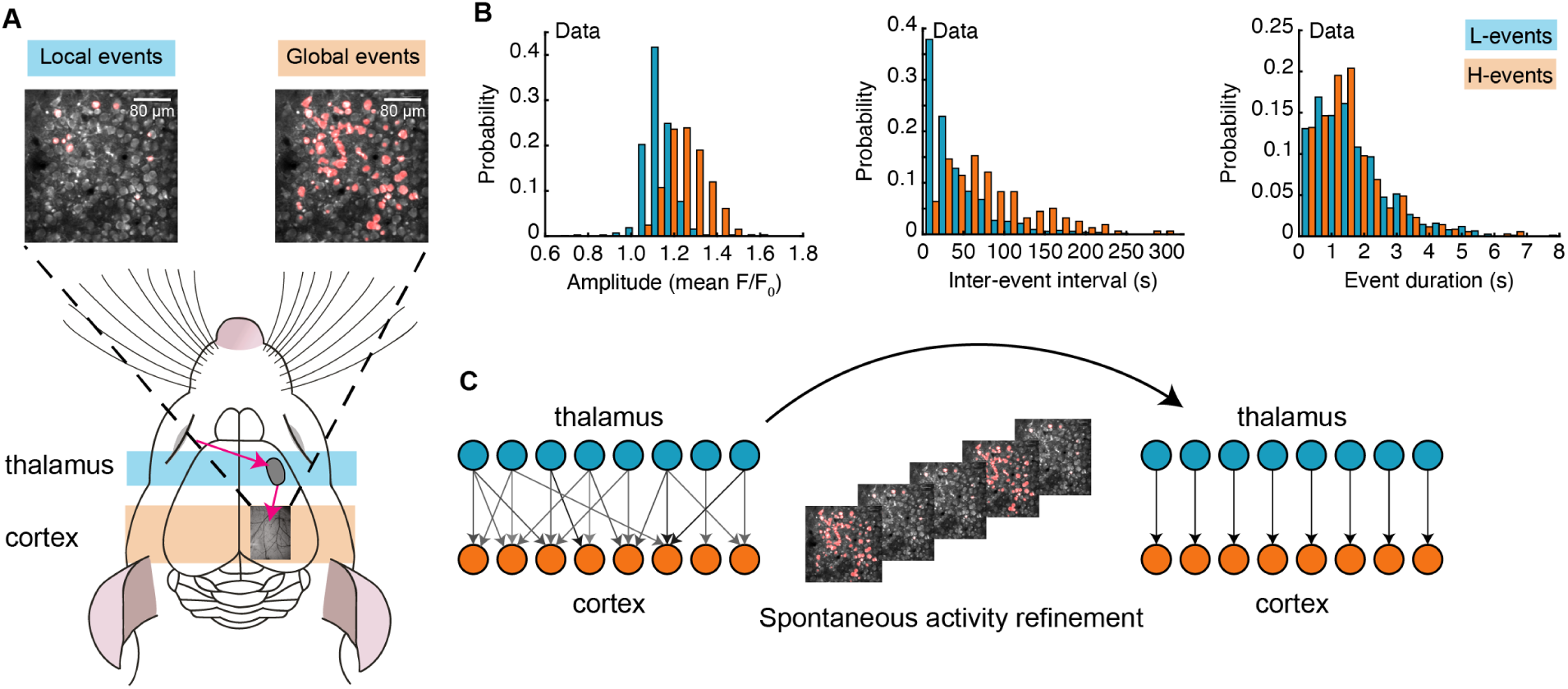
Spontaneous activity patterns in early postnatal development. **A**. Two distinct patterns of spontaneous activity recorded *in vivo* in the visual cortex of young mice before eye-opening (P8-10). Blue shading denotes local spontaneous events (L-events; generated by retinal waves); orange shading denotes global spontaneous events (H-events). Activated neurons during each event are shown in red. **B**. Distributions of different event properties (amplitude, inter-event interval and event duration). Amplitude was measured as changes in fluorescence, relative to baseline, F/F0. **C**. Network schematic: thalamocortical connections are refined by spontaneous activity. The initially broad receptive fields with weak synapses evolve into a stable configuration with strong synapses organized topographically.

Next, we built a model that incorporates these two different patterns of spontaneous activity to investigate how they jointly refine network connectivity between the thalamus and the visual cortex (Fig. 1C). We used a one-dimensional feedforward network model – a microcircuit motivated by the small region of cortex imaged experimentally – composed of two layers, an input (presynaptic) layer corresponding to the sensory periphery (the thalamus) and a target (postsynaptic) layer corresponding to the primary visual cortex (Fig. 2A). Cortical activity ***v*** in the model is generated by two sources (Fig. 2B; Table 1): First, by L-events, **u**, which activate neighboring thalamic cells and drive the cortex through the weight matrix, ***W***, and second, by H-events, ***v***^spon^, activating the majority of the cortical cells. We used a rate-based unit with a membrane time constant *τ*_*m*_ and linear activation function consistent with the coarse temporal structure of spontaneous activity during development, carrying information on the order of hundreds of milliseconds [13, 36, 3]:

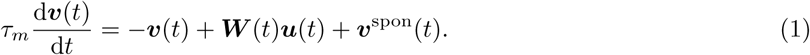

**Table 1.** List of parameters used in the model unless stated otherwise.

| Name | Value/Distribution | Description |
| --- | --- | --- |
| **Network** |  |  |
| $N_u$ | 50 | number of thalamic neurons |
| $N_v$ | 50 | number of cortical neurons |
| $T$ | 50000 | simulation length [s] |
| **Weights** |  |  |
| $w_{\text{ini}}$ | U(0.15,0.25) | range of initial weights (U: uniform dist.) |
| $s$ | 0.05 | amplitude of Gaussian bias |
| $\sigma_s$ | 4 | spread of Gaussian bias |
| $w_{\text{max}}$ | 0.5 | weight saturation limit |
| **L-events** |  |  |
| $L_{\text{amp}}$ | 1.0 | amplitude (equivalently, binary neuron) |
| $L_{\text{pct}}$ | U(20%,80%) | percentage of thalamic cells activated |
| $L_{\text{dur}}$ | $\mathcal{N}(0.15, 0.015)$ | mean duration [s] ($\mathcal{N}$: Gaussian dist.) |
| $L_{\text{int}}$ | Exp(1.5) | mean inter-event interval [s] (Exp: exponential dist.) |
| **H-events** |  |  |
| $H_{\text{amp}}$ | $\mathcal{N}(6, 2)$ | amplitude |
| $H_{\text{pct}}$ | U(80%,100%) | percentage of cortical cells activated |
| $H_{\text{dur}}$ | $\mathcal{N}(0.15, 0.015)$ | mean duration [s] |
| $H_{\text{int}}$ | Gamma(3.5, 1.0) | mean inter-event interval [s] (Gamma: Gamma dist.) |
| **Time constants** |  |  |
| $\tau_m$ | 0.01 | membrane time constant [s] |
| $\tau_w$ | 500 | weight-change time constant for Hebbian rule [s] |
| $\tau_w$ | 1000 | weight-change time constant for BCM rule [s] |
| $\tau_\theta$ | 20 | output threshold time constant for BCM rule [s] |
| $\tau_\eta$ | 1 | adaptation time constant [s] |

**Figure 2.**
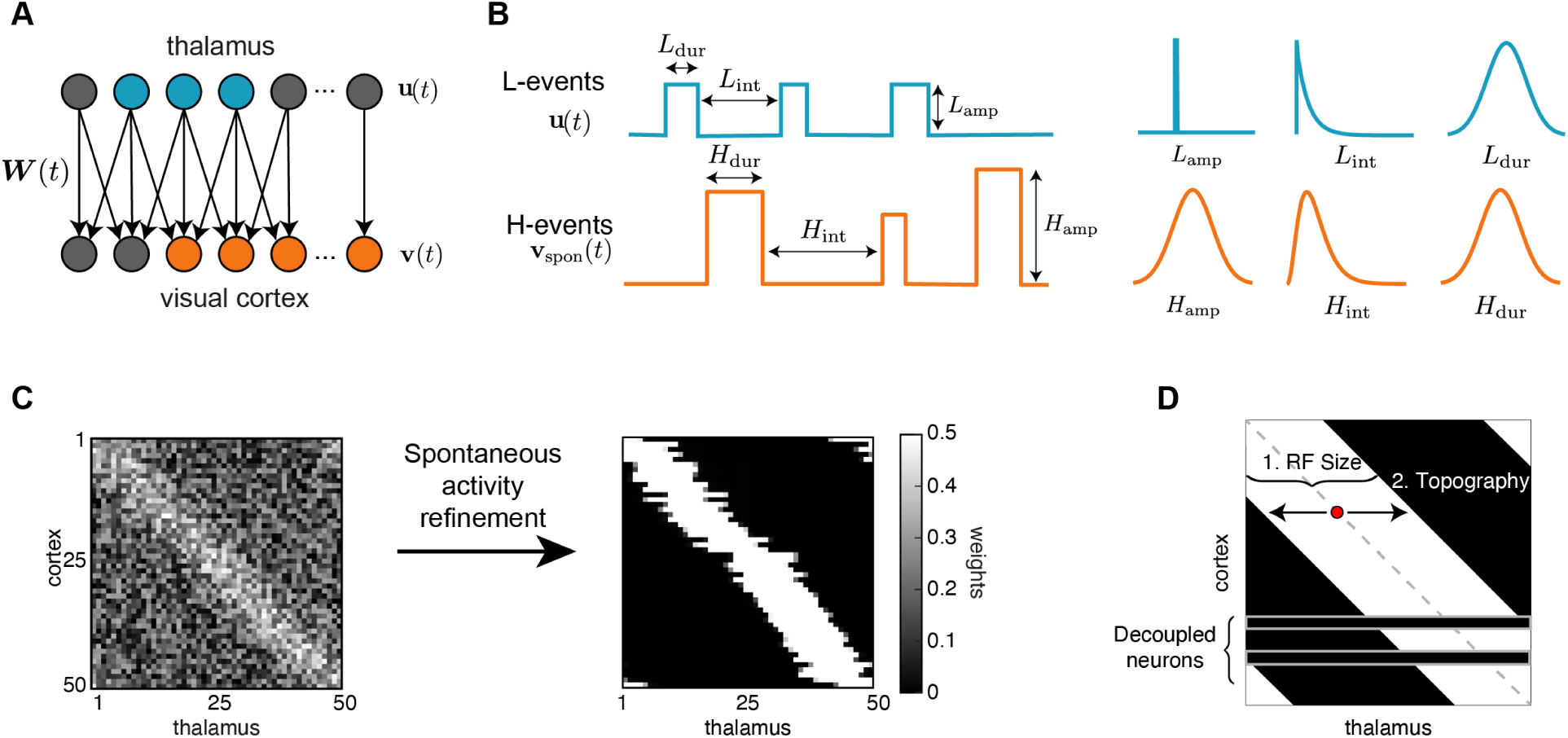
A network model of thalamocortical connectivity refinements. **A**. A feedforward network with an input layer of thalamic neurons ***u***(*t*) connected to an output layer of cortical neurons ***v***(*t*) by synaptic weights ***W*** (*t*). **B**. Properties of L- and H-events in the model (amplitude *L*_amp_, *H*_amp_, inter-event interval *L*_int_, *H*_int_ and duration *L*_dur_, *H*_dur_) follow probability distributions extracted from data [26] (see Table 1). **C**. Initially weak all-to-all connectivity with a small bias along the diagonal (left), corresponding to perfect topography, gets refined by the spontaneous activity events (right). **D**. Evaluating network refinement through receptive field statistics (see Methods). We quantify two properties: 1. the receptive field size and 2. the topography, which quantifies on average how far the receptive field center of each cortical cell (red dot) is from the diagonal (dashed gray line). We also determined how many cortical cells decouple from the thalamus (see Methods).

To investigate the refinement of network connectivity during development, we studied the evolution of synaptic weights using rate-based plasticity rules operating over long timescales identified experimentally [37, 38]. First, we examined a classical Hebbian plasticity rule where coincident presynaptic thalamic activity and postsynaptic cortical activity in the form of L-events leads to synaptic potentiation. We postulated that H-events act homeostatically, maintaining synaptic weights in an operating regime by triggering global network depression involving the majority of synaptic weights in the absence of peripheral drive. Because they activate many neurons simultaneously, H-events lack the potential to drive topographical refinements. Their postulated homeostatic action resembles synaptic depression through downscaling, as observed in response to highly-correlated network activity, for instance upon blocking inhibition [39], or during slow-wave sleep [40]. Therefore, we added a non-Hebbian term that depends only on the postsynaptic activity, with a proportionality constant which controls the relative amount of synaptic potentiation vs. depression. This differs from other Hebbian covariance plasticity rules for the generation of weight selectivity, which include non-Hebbian terms that depend on both pre- and postsynaptic activity [31, 41] and is mathematically related to models of heterosynaptic plasticity [42, 43, 44]. In this Hebbian rule, the change in synaptic weight between cortical neuron *j* and thalamic neuron *i* is given by:

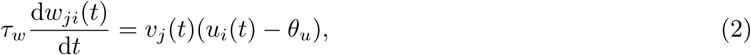

where *τ*_*w*_ is the learning time constant and *θ*_*u*_ the proportionality constant in the non-Hebbian term, which we refer to as the ‘input threshold’. The activity time constant *τ*_*m*_ is much faster than the learning time constant, *τ*_*m*_ ≪ *τ*_*w*_, which allows us to separate timescales and to study how network activity on average affects learning (see Appendix 1).

Next, we investigated the BCM learning rule, commonly used in theoretical studies for its ability to generate selectivity in postsynaptic neurons which experience patterned inputs [32]. An important property of the BCM rule is its ability to homeostatically regulate the balance between potentiation and depression of all incoming inputs into a given neuron depending on the difference between the activity of that neuron and some target level. The change in synaptic weight between cortical neuron *j* and thalamic neuron *i* is given by:

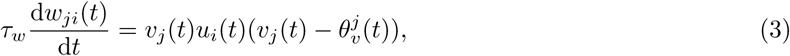

where

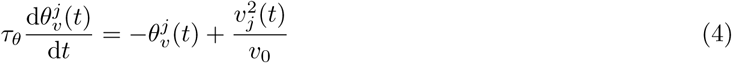

describes how the sliding threshold 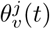 depends on postsynaptic activity, *v*_0_ is the target rate of the cortical neurons and *τ*_*θ*_ the sliding threshold time constant. According to this rule, synaptic weight change is Hebbian in that it requires coincident pre- and postsynaptic activity, as is only the case during L-events. H-events induce no direct plasticity in the network because of the absence of presynaptic activation, but they still trigger synaptic depression indirectly by increasing the sliding threshold between potentiation and depression.

Based on experimental measurements of the extent of thalamocortical connectivity at different developmental ages [45], we assumed that initial network connectivity was weak and all-to-all, such that each cortical neuron was innervated by all thalamic neurons. To account for the activity-independent stage of development guided by molecular guidance cues and chemoaffinity gradients [1, 46], a small bias was introduced to the initial weight matrix to generate a coarse topography in the network, where neighboring neurons in the thalamus project to neighboring neurons in the cortex and preserve spatial relationships (Fig. 2C, left). Following connectivity refinements through spontaneous activity and plasticity, a desired outcome is that the network achieves a stable topographic configuration (Fig. 2C, right) where each cortical neuron receives input only from a neighborhood of thalamic neurons. To evaluate the success of this process, we quantified two properties. First, the receptive field size defined as the average number of thalamic neurons that strongly innervate a cortical cell (Fig. 2D, 1). We normalized the receptive field size to the total number of thalamic cells, so that it ranges from 0 (no receptive field, all cortical cells decouple from the thalamus) to 1 (each cortical cell receives input from all the thalamic cells, all weights potentiate leading to no selectivity). We also quantified the topography of the final receptive field (Fig. 2D, 2 and Methods), which evaluates how well the initial bias is preserved in the final network connectivity. The topography ranges from 0 (all cortical neurons connect to the same set of thalamic inputs) to 1 (perfect topography relative to the initial bias). We note that the lack of initial connectivity bias did not disrupt connectivity refinements and receptive field formation, but could not on its own establish topography (Fig. S S1A). We also observed that in many cases cortical cells decouple from the thalamus.

### Spontaneous cortical H-events disrupt topographic connectivity refinement

Both the Hebbian and the BCM learning rules are known to generate selectivity with patterned input stimuli [41, 32], and we confirmed that L-events can refine receptive fields in both scenarios (Fig. S1B, C). We found that including H-events in the Hebbian rule requires that the input threshold *θ*_*u*_ and the inter-event interval for H-events *H*_int_ be precisely balanced to generate selectivity and appropriately refined receptive fields (Fig. 3A,C-left). For a narrow range of *H*_int_, weight selectivity emerges, but with some degree of decoupling between pre- and postsynaptic neurons (Fig. 3A, middle). Outside of this narrow functional range, individual cortical neurons are either non-selective (Fig. 3A, left) or decoupled from the thalamus (Fig. 3A, right). In comparison, including H-events in the BCM learning rule does not decouple pre- and postsynaptic neurons (Fig. 3B and Fig. S2) and selectivity can be generated over a wider range of H-inter-event-intervals *H*_int_ and target rates *v*_0_ for the BCM rule (Fig. 3C, right).

**Figure 3.**
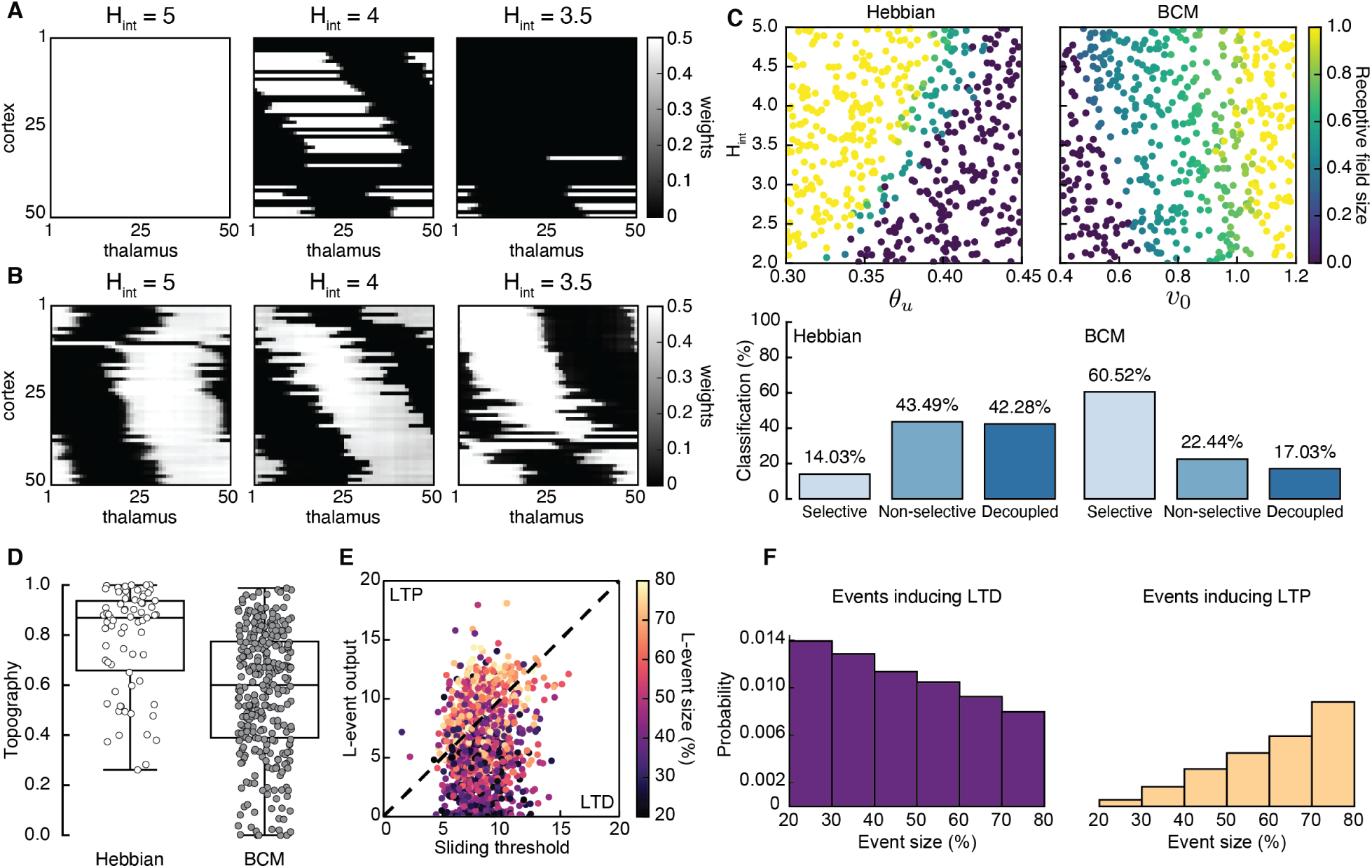
Spontaneous cortical events disrupt receptive field refinement. **A**. Receptive fields generated by the Hebbian rule with input threshold *θ*_*u*_ = 0.4 and increasing *H*_int_. **B**. Receptive fields generated by the BCM rule with target rate *v*_0_ = 0.7 and increasing *H*_int_. **C**. Top: Average receptive field sizes obtained from 500 Monte Carlo simulations for combinations of *H*_int_ and *θ*_*u*_ for the Hebbian rule (left) and *H*_int_ and *v*_0_ for the BCM rule (right). Bottom: Percentage of simulation outcomes classified as ‘selective’ when the average receptive field size is smaller than 1 and larger than 0, ‘non-selective’ when the average receptive field size is equal to 1, and ‘decoupled’ when the average receptive field size is 0 for the two rules. **D**. Topography of receptive fields classified as selective in C. Topography for the Hebbian rule: 0.78 ± 0.20, and for the BCM rule: 0.57 ± 0.25 (mean ± standard deviation). **E**. The response of a single cortical cell to L-events of different sizes (color) as a function of the sliding threshold for the BCM rule with *H*_int_ = 3.5 and *v*_0_ = 0.7. The cell’s incoming synaptic weights from presynaptic thalamic neurons undergo LTP or LTD depending on L-event size. **F**. Probability of L-event size contributing to LTD (left) and LTP (right) for the BCM rule with the same parameters as in E.

Despite this apparent advantage of the BCM rule, it generates receptive fields with much worse topography than the Hebbian learning rule (Fig. 3D). We found that the underlying reason for this worse topography of the BCM rule is the sign of synaptic change evoked by L-events of different sizes. In particular, small L-events generate postsynaptic cortical activity smaller than the sliding threshold and promote long-term synaptic depression (LTD), while large L-events generate cortical activity larger than the sliding threshold and promote long-term synaptic potentiation (LTP) (Fig. 3E,F). Therefore, the amount of information for connectivity refinements present in the small L-events is limited in the BCM learning rule resulting in poor topographic organization of receptive fields.

Taken together, our results confirm that H-events can operate in parallel to network refinements by L-events homeostatically regulating connection strength as postulated. However, the formation of receptive fields in the Hebbian rule is very sensitive to small changes in event properties (e.g., inter-event intervals), which are common throughout development [33]. In this case, H-events are disruptive and lead to the elimination of all thalamocortical synapses, effectively decoupling the cortex from the sensory periphery. In the BCM rule, including H-events prevents the decoupling of cortical cells from the periphery because the amount of LTD is dynamically regulated by the sliding threshold on cortical activity. However, L-events lose the ability to instruct topography because they generate LTP primarily when they are large.

### Adaptive H-events achieve robust selectivity

Upon comparing the distinct outcomes of the Hebbian and BCM learning rules, we proposed that a mechanism that regulates the amount of LTD during H-events based on cortical activity, similar to the sliding threshold of the BCM rule, could be a potential biologically plausible solution to mitigate the decoupling of cortical cells in the Hebbian rule. Therefore, this mechanism combined with the Hebbian learning rule could lead to refined receptive fields that also have good topographic organization. Hence, we postulated that H-events adapt by assuming that during H-events cortical cells scale their amplitude to the average amplitude of the preceding recent events. In particular, for each cortical cell *j* an activity trace *η*_*j*_ integrates the cell’s firing rate *v*_*j*_ over a timescale *τ*_*η*_ slower than the membrane time constant:

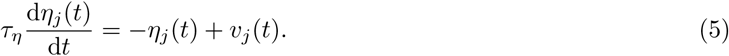

This activity trace *η*_*j*_ then scales the intrinsic firing rate of the cortical cells during an H-event, *H*_amp_ *→ η*_*j*_*H*_amp_, making it dependent on its recent activity. The activity trace *η*_*j*_ might biophysically be implemented through a calcium-dependent signaling pathway that is activated upon sufficient burst depolarization and that is able to modulate a cell’s excitability in the form of intrinsic plasticity [47, 48, 49]. A fast, activity-dependent mechanism that decreases single-neuron excitability following a prolonged period of high network activity has been identified in spinal motor neurons of neonatal mice [50]. However, there might be other ways to implement this adaptation (see Discussion).

Using adaptive H-events, we investigated the refinement of receptive fields in the network with the same Hebbian rule (Fig. 4A). In sharp contrast to the Hebbian rule with non-adaptive H-events (Fig. 3A), we observed that changing the average inter-event interval of H-events in a wider and more biologically realistic range (from the data, *H*_int_ *∼* 3*L*_int_) yields selectivity and appropriately refined receptive fields (Fig. 4A). Increasing *θ*_*u*_ or decreasing *H*_int_ yields progressively smaller receptive fields, while mitigating cortical decoupling (Fig. 4B). The refined receptive fields also have a very good topography because L-events in the Hebbian learning rule carry nearest-neighbor information for the topographic refinements (Fig. 4C). Next, we investigated how the proposed adaptive mechanism scales H-event amplitude by modulating the relative strength of H-events. For the Hebbian rule, we calculated the analytical solution for weight development with L- and H-events by reducing the dimension of the system to two: one being the average of the weights that potentiate and form the receptive field, *w*_RF_, and the other being the average of the remaining weights, which we called ‘complementary’ to the receptive field, *w*_C_ (Fig. 4D; see Appendix 1, Methods). We calculated the phase plane area of the reduced two-dimensional system with non-adaptive H-events (calculated as the proportion of initial conditions) that results in selectivity, potentiation or depression (Fig. 4D, bottom). We found that adaptively modulating the strength of H-events maximizes the area of the phase plane that results in selectivity (Fig. 4E, shaded region). The range of H-event strengths that maximizes the selective area for each input threshold in the reduced two-dimensional system can be related to the scaling of H-event amplitude in the simulations (Methods). In particular, the adaptation reliably shifts the H-event amplitude that would have occurred without adaptation, ‘non-adapted strength of H-events’, into the regime of amplitudes that maximizes selectivity, ‘adapted strength of H-events’ (Fig. 4E). Therefore, the adaptation of H-event amplitudes controls the selective refinement by peripheral L-events by modulating cortical depression by the adapted H-events.

**Figure 4.**
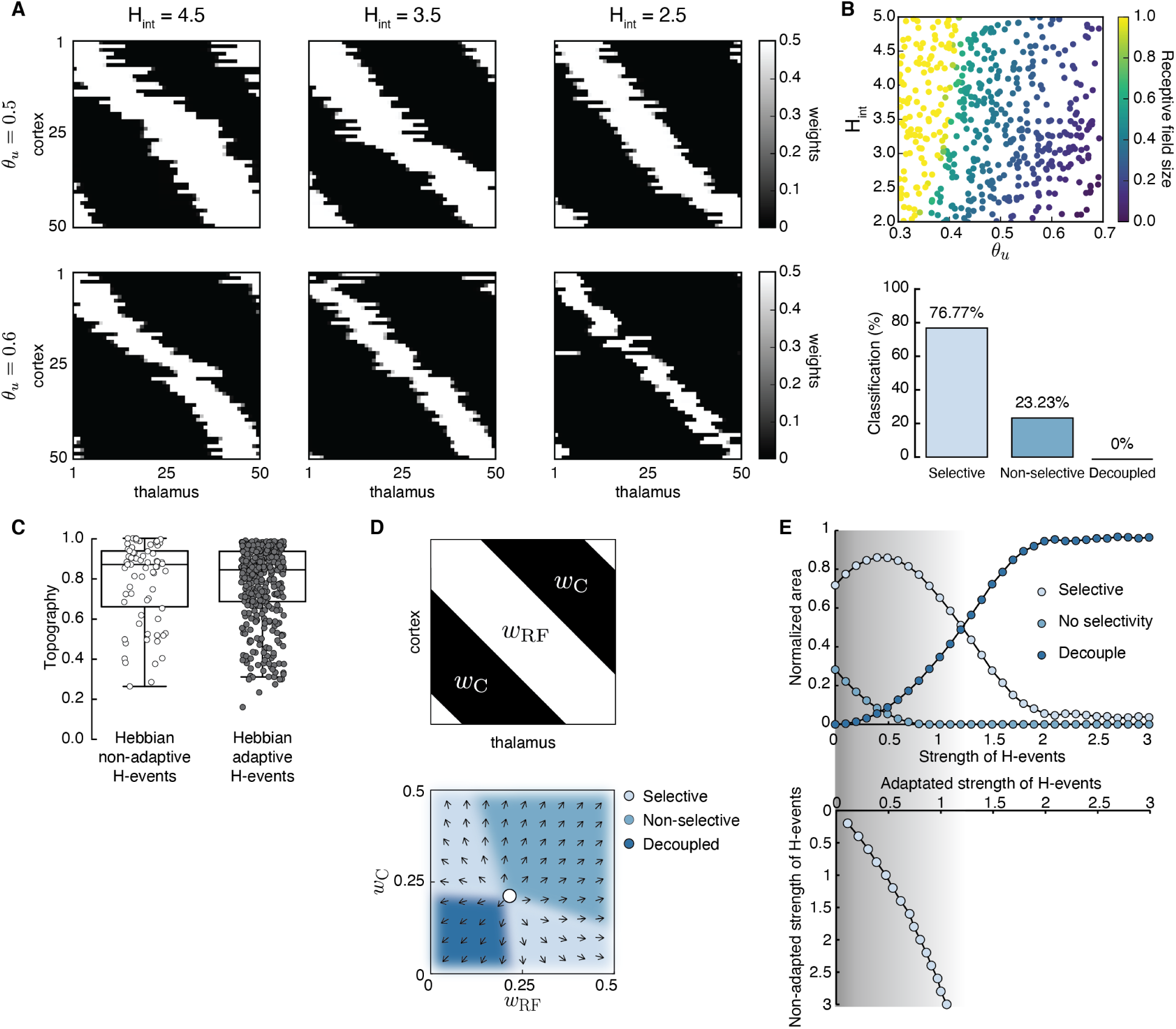
Adaptive cortical events refine thalamocortical connectivity. **A**. Receptive field refinement with adaptive H-events and different H-inter-event-intervals, *H*_int_. Top: *θ*_*u*_ = 0.5; bottom: *θ*_*u*_ = 0.6. **B**. Receptive field sizes from 500 Monte Carlo simulations for combinations of *H*_int_ and *θ*_*u*_. Bottom: Percentage of simulation outcomes classified as ‘selective’ when the average receptive field size is smaller than 1 and larger than 0, ‘non-selective’ when the average receptive field size is equal to 1, and ‘decoupled’ when the average receptive field size is 0 for the two rules. **C**. Topography of receptive fields classified as selective in B using non-adaptive (0.78±0.20) and adaptive (0.78±0.19) H-events (mean ± standard deviation). **D**. Top: Reduction of the full weight dynamics into two dimensions. Two sets of weights were averaged: those which potentiate and form the receptive field, *w*_RF_, and the complementary set of weights that depress, *w*_C_. Bottom: Initial conditions in the reduced two-dimensional phase plane were classified into three outcomes: ‘selective’, ‘non-selective’ and ‘decoupled’. We sampled 2500 initial conditions which evolved according to Eq. A4 until the trajectories reached one of the selective fixed points, (*w*_max_, 0) and (0, *w*_max_), or resulted in no selectivity either because both weights depressed to (0, 0) or potentiated to (*w*_max_, *w*_max_). The normalized number of initial coordinates generating each region can be interpreted as the area of the phase plane that results in each outcome. **E**. Top: Normalized area of the phase plane of the reduced two-dimensional system that resulted in ‘selective’, ‘non-selective’ and ‘decoupled’ outcomes for *θ*_*u*_ = 0.53 as a function of H-events strength. The darker shading indicates ranges of non-adapted H-event strength where the selectivity area is maximized. Bottom: The corresponding adapted strength of H-events was calculated in simulations with adaptive H-events, and plotted as a function of the nominal, non-adaptive strength of H-events. The range of adapted H-event strengths (bottom) corresponds to the range of non-adaptive values that maximize the selectivity area (top). Each point shows the average over 10 runs and the bars the standard deviation (which are very small).

### *In vivo* spontaneous cortical activity shows a signature of adaptation

To determine whether spontaneous cortical activity contains a signature of our postulated adaptation mechanism of H-event amplitudes, we reanalyzed published *in vivo* two-photon Ca^2+^ imaging data recorded in the visual cortex of young mice (P8-10) [26]. We found no evidence for long-term fluctuations in cortical excitability, making it unlikely that L- and H-event amplitudes are co-regulated by activity-independent mechanisms (Fig. S3). We identified L- and H-events based on previously established criteria, including participation rates, event amplitude and jitter [26]. For each detected H-event, we extracted all spontaneous (L- or H-) events that preceded this H-event combining several consecutive recordings in the same animal, up to a maximum time *τ*_max_ = 300 s. We then scaled the amplitude of each previous event by an exponential kernel with a decay time constant of *τ*_decay_ = 1000 s, which is sufficiently long to integrate many preceding spontaneous events (compared with the inter-event intervals in Fig. 1B), and averaged these scaled amplitudes to get an aggregate quantity over amplitude and frequency (see Methods).

We found that this aggregate amplitude of L- and H-events preceding a given H-event is significantly correlated (*r* = 0.44, *p* < 10^−10^) to the amplitude of the selected H-event (Fig. 5B). Consequently, a strong (weak) H-event follows strong (weak) average preceding network activity (Fig. 5C), suggesting that cortical cells indeed adapt their spontaneous firing rates as a function of their previous activity levels. The correlations are robust to variations in the inclusion criteria, maximum time *τ*_max_ to integrate activity and the exponential decay time constant *τ*_decay_ (Fig. S4). This analysis suggests that the timescale of adaptation is reasonably fast, but at the same time sufficiently long to integrate preceding spontaneous events, matching the relevant timescales of information transmission in the developing cortex [51, 13].

**Figure 5.**
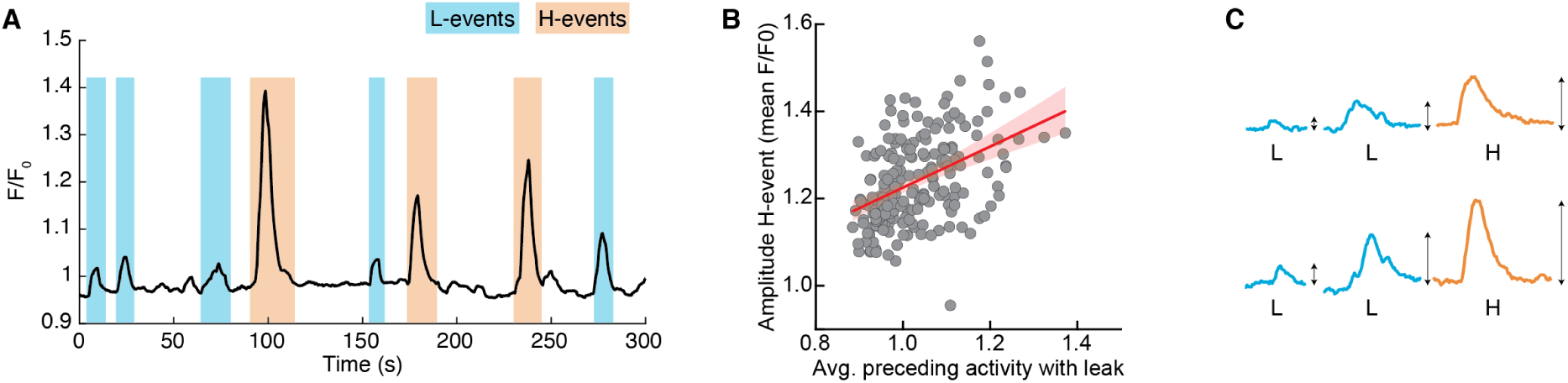
Spontaneous events in developing cortex adapt to recent activity. **A**. Calcium trace of a representative recording with L- (blue) and H-events (orange) [26]. For each H-event, the amplitude of the spontaneous events (both L- and H-) that preceded it within a time window *τ*_max_ = 300 s was averaged with an exponential leak of decay time constant *τ*_decay_ = 1000 s. **B**. Significant correlation between the amplitude of an H-event and the average amplitude of preceding events within *τ*_max_ (*N* = 195 events from 9 animals). Animals with fewer than 12 H-events preceded by activity within *τ*_max_ were excluded from this analysis (see Methods). The Pearson correlation coefficient is *r* = 0.44 *p* < 10^−10^, CI = (0.32, 0.54). Red line indicates regression line with 95% confidence bounds as shaded area. **C**. Schematic of the postulated adaptation: A weak (strong) H-event is more likely to be preceded by weak (strong) spontaneous events.

### Modulating spontaneous activity properties makes different predictions for receptive field refinements

Our results make relevant predictions for the refinement of receptive fields upon manipulating spontaneous activity. H-event frequency can be experimentally reduced by a gap junction blocker (carbenoxolone) [26]. L-events can also be experimentally manipulated, for instance, by altering inhibitory signaling [35], or the properties of retinal waves which propagate as L-events into the cortex. While the effect of altered inhibitory signaling on receptive field refinements is still unknown, retinal wave manipulations evoke measurable defects in the retinotopic map refinement of downstream targets [52, 53, 12]. A most prominent example are *β*2 knockout mice, which lack expression of the *β*2 subunit of the nicotinic acetylcholine receptor (*β*2-nAChR) that mediates spontaneous retinal waves in the first postnatal week. In these animals, retinal waves are consistently larger as characterized by the increased correlation with distance [54, 55, 56], in addition to other features. As a result, the receptive fields in the visual cortex in *β*2-mutants are imprecise and less refined. Based on the analysis of our model (Fig. 4), we asked what implications the spatial structure of L-events might have on receptive field refinement.

We performed Monte Carlo simulations with a range of input thresholds *θ*_*u*_ and variable participation rates of thalamic neurons in L-events, using the Hebbian rule with adaptive H-events (Fig. 6A). Consistent with the imprecise receptive fields in the visual cortex in *β*2-mutants [53], larger L-events in our model produce less refined i.e. larger receptive fields in the cortical network (Fig. 6B, C left). Smaller L-events also refine receptive fields with better topographic organization (Fig. 6C, right) and do not impair connectivity refinements. This result could be linked to experiments where the expression of *β*2-nAChR is limited to the ganglion cell layer of the retina, resulting in smaller retinal waves than those in wild-type and undisturbed retinotopy in the superior colliculus [11], although the effects in the cortex are unknown. Therefore, we propose that manipulations that modulate the size of sensory activity from the periphery have a profound impact on the precision of receptive field refinement in downstream targets, making predictions to be tested experimentally.

**Figure 6.**
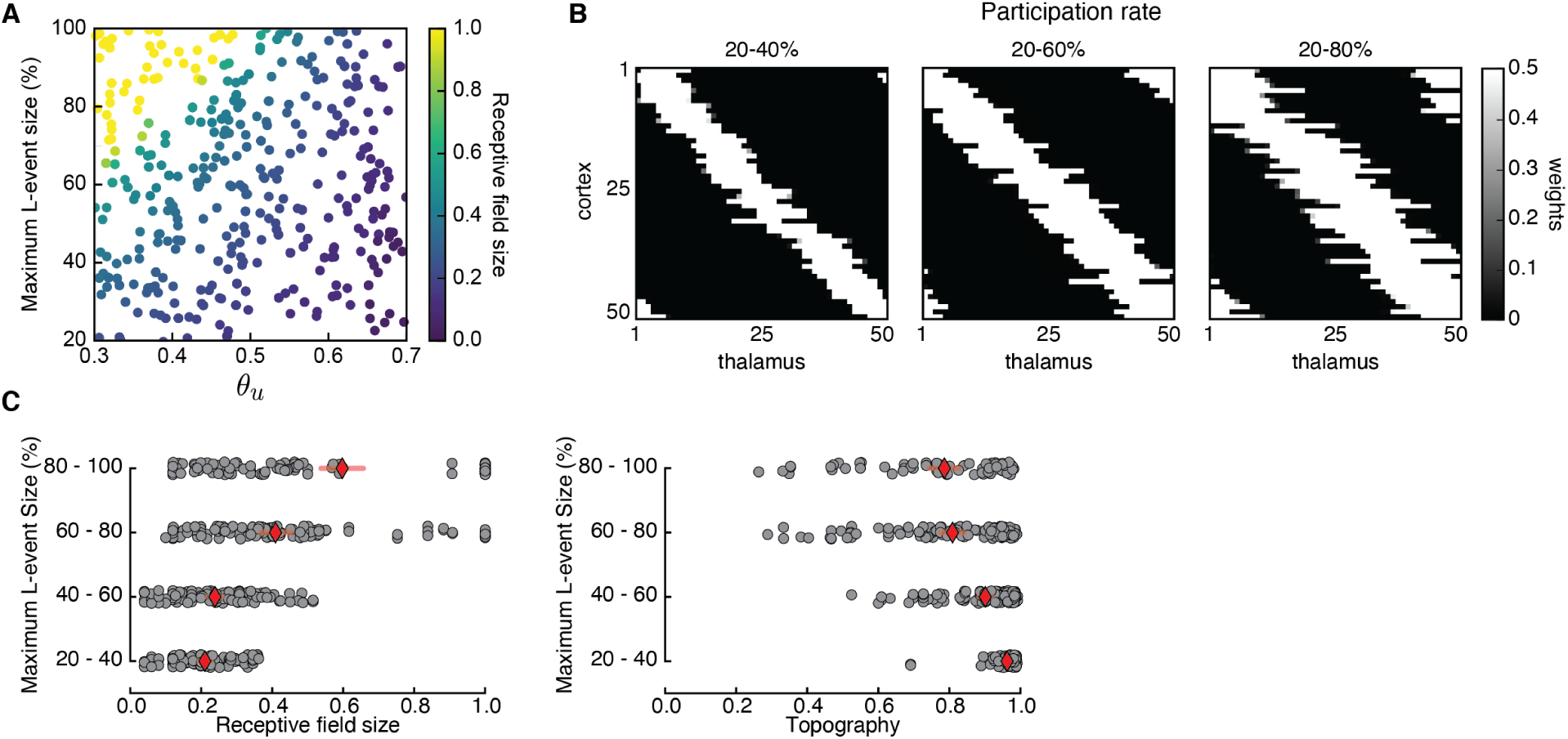
Receptive field refinement depends on the properties of L-events. **A**. Receptive field sizes from 500 Monte Carlo simulations for different sizes of L-events where the minimum participation rate was 20% and the maximum participation rate was varied. The input threshold was taken from the range 0.3 ≤ *θ*_*u*_ ≤ 0.7, while the adaptive H-events had a fixed inter-event-interval *H*_int_ = 3.5. **B**. Individual receptive fields for different L-event maximum participation rates and *θ*_*u*_ = 0.50. As the upper bound of the participation rate progressively increases from 40% to 80%, receptive fields get larger. **C**. Left: Receptive field sizes from A binned according to the maximum L-event size. Right: Corresponding topography of selective receptive fields for different sizes of L-events.

### Adaptive H-events promote the developmental event sparsification of cortical activity

Spontaneous activity is not static, but consists of distinct patterns that are dynamically regulated during development by ongoing activity-dependent plasticity which continuously reshapes network connectivity [33, 34, 57]. We next asked how our observed modifications in network connectivity that result in refined receptive fields based on fixed activity statistics during a single postnatal day might further modify spontaneous activity patterns on a much longer developmental timescale of several days. Therefore, we analyzed the activity of the simulated cortical neurons during the process of receptive field refinement in the presence of adaptive H-events. We detected ‘effective’ L- and H-events in the simulated cortical activity following an automatic event detection similar to the routine used to detect events in the experimental calcium traces (Figures 7A and B, see Methods for details). Given the sequence of these simulated effective L- and H-events, we reconstructed the probability distributions of their amplitudes, inter-event intervals and durations as in the data (Fig. 7B and Fig. S5A). Here we stress that the features of effective events differ from the features of ‘putative’ ones that were initially modeled according to the data (Table 1). During a putative H-event, the adaptation mechanism can decrease the amplitude of simulated cortical cells. Upon sufficient reduction in amplitude, such a putative H-event can have a reduced effective participation rate because only the subset of the cortical neurons that is activated above the detection threshold will be detected as active. Therefore, a putative H-event can be identified as an effective L-event.

**Figure 7.**
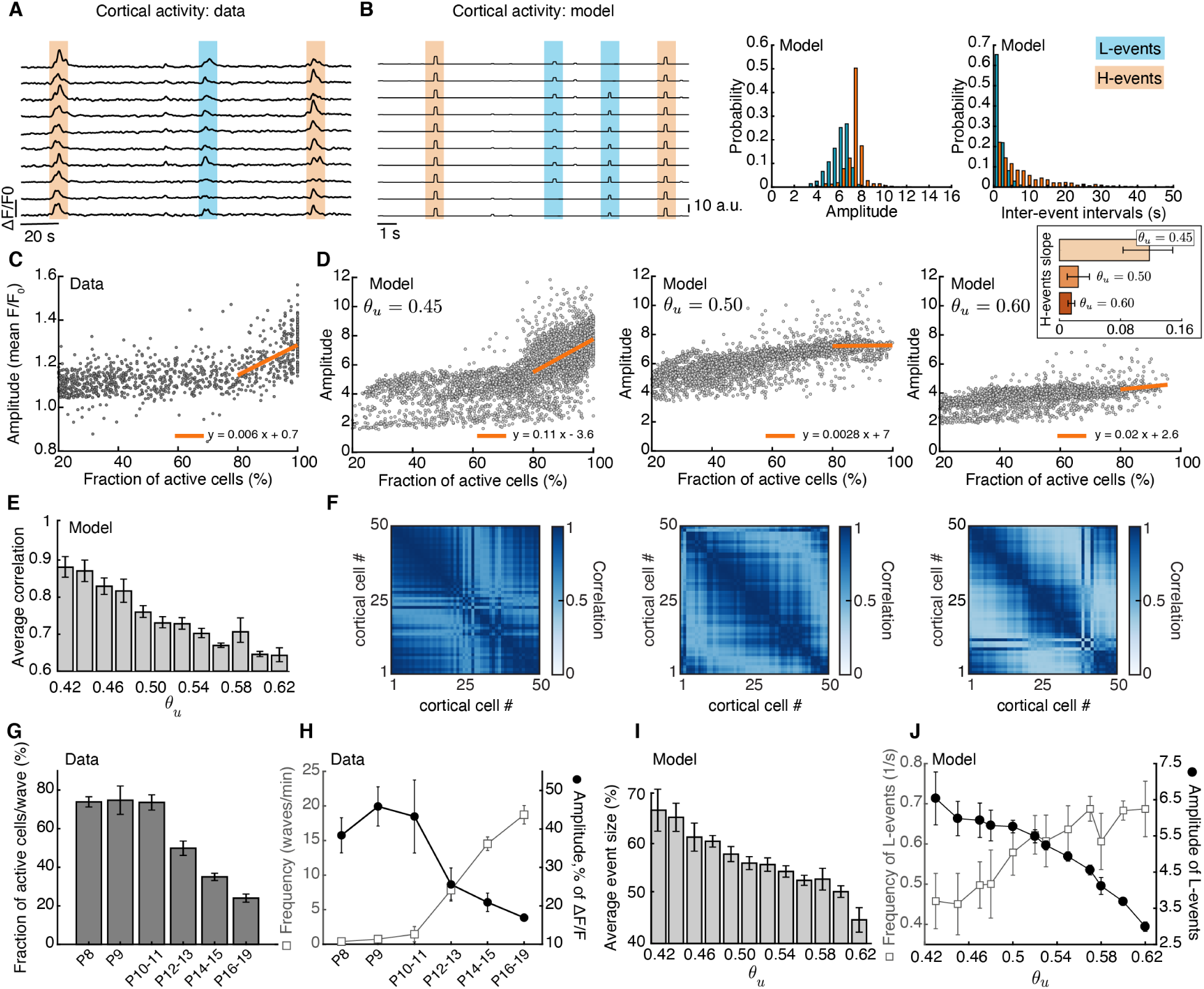
Adaptive H-events promote sparsification of cortical activity during development. **A**. Identifying L- and H-events in mouse cortical activity [26]. Each activity trace represents an individual cortical cell. **B**. Left: We used a similar protocol to identify effective L- and H-events in the model. Right: Distributions of average amplitude and inter-event intervals extracted from the model. **C**. Amplitude *v* s. participation rate plot from the data [26]. The regression line for the amplitude vs. participation rate in H-events has a positive slope. **D**. Amplitude *v* s. participation rate plots from the model, for different values of *θ*_*u*_. Inset: The regression line for the amplitude vs. participation rate in H-events has a slope that decreases with *θ*_*u*_. **E**. Correlation between cortical neurons decreases as a function of the input threshold *θ*_*u*_ in the model, a proxy for developmental time. **F**. Correlation matrices of simulated cortical neuron activity corresponding to D. **G,H**. Event sizes and the relationship between frequencies (open squares) and amplitudes (filled circles) of spontaneous events at different postnatal ages (data reproduced from [33]). **I**. Effective spontaneous event sizes as a function of the input threshold *θ*_*u*_. **J**. Frequencies (squares) and amplitudes (circles) of effective L-events in the model at different input thresholds. For the data (model), the bars represent the standard error of the mean (standard deviation of 10 simulation runs)

What is the effect of this classification of putative H-events as effective L-events? The amount of putative H-events classified as effective L-events progressively increases as receptive fields are further refined, which represents developmental time in the model. Since the input threshold *θ*_*u*_ of the Hebbian learning rule is clearly related to receptive field size (Fig. 4B), we used *θ*_*u*_ as a proxy for time of development in the model: low *θ*_*u*_ corresponds to earlier developmental stages when receptive fields are large, while high *θ*_*u*_ corresponds to late developmental stages when receptive fields are refined. This assumption is also in line with the fact that the input resistance of neurons in V1 and S1 decreases during development [58, 34], so that the depolarizing current necessary to trigger an action potential increases with age.

We investigated the progressive evolution of spontaneous activity properties in our network model by varying *θ*_*u*_. At an early developmental stage in the model (*θ*_*u*_ = 0.45), the relationship between effective event amplitude and participation rate (Fig. 7D, left) strongly resembles the data (Fig. 7C, reproduced from [26]). The amplitude of effective L-events is approximately half the amplitude of effective H-events. Moreover, we also observe a high proportion of events with large sizes (Fig. S5B), suggesting that in this network large spontaneous events are very frequent. At an intermediate developmental stage in the model (*θ*_*u*_ = 0.50), the resemblance between effective amplitude and participation rate to the data is less pronounced (Fig. 7D, middle), with lower average amplitudes and density of large events (Fig. S5B). Finally, at late developmental stages in the model (*θ*_*u*_ = 0.60), the dependence of effective amplitude on participation rate is almost absent (Fig. 7D, right). Event amplitude is much lower and there are fewer effective events with more than 80% participation rate (Fig. S5C). Therefore, we observe a progressive event sparsification of the effective spontaneous events during ongoing development in our model, whereby spontaneous events become smaller in size, with fewer cells participating. This event sparsification also becomes apparent when examining the correlation matrices of simulated cortical activity for increasing input threshold values (Fig. 7E, F).

Interestingly, such event sparsification of spontaneous activity has been observed experimentally in the mouse barrel cortex during postnatal development from P4 to P26 [34] and in the visual cortex from P8 to P79 [33]. During this period, in the visual cortex, the size of spontaneous events decreases (Fig. 7G), the amplitude of the participating cells also decreases, while event frequency increases (Fig. 7H) [33]. This progressive event sparsification of cortical activity is generated by mechanisms intrinsic to the cortex, and does not seem to be sensory-driven [33, 34]. We found the same relationships in our model using *θ*_*u*_ as a proxy for developmental time (Fig. 7I, J).

In summary, our framework for activity-dependent plasticity and receptive field refinement between thalamus and cortex with adaptive H-events can tune the properties of cortical spontaneous activity and provide a substrate for the event sparsification of cortical activity during development on a much longer timescale than receptive field refinement. This sparsification has been found in different sensory cortices, including visual [33], somatosensory [34] and auditory [57], suggesting a general principle that underlies network refinement. However, the event sparsification we observe is different than sparse network activity implicated in sparse efficient coding, which interestingly seems to decrease during development [59, 60]. Our modeling predicts that cortical event sparsification is primarily due to the suppression of cortically-generated H-events in the Hebbian rule, which switches cortical sensitivity to input from the sensory periphery after the onset of sensory experience.

## Discussion

We examined the information content of spontaneous activity for refining local microcircuit connectivity during early development. In contrast to classical works on activity-dependent refinements which used mathematically convenient formulations of spontaneous activity [61, 41], we used spontaneous activity patterns characterized in the mouse visual cortex *in vivo* before the onset of vision, which revealed its rich structure. Specifically, we explored the joint contribution of two distinct patterns of spontaneous activity recorded in the mouse visual cortex before the onset of vision, local (L-events) and global (H-events), on robustly establishing precise topographic receptive fields between the thalamus and the cortex in a rate-based network model with activity-dependent plasticity. We compared the emergence of selective and topographic receptive fields using two learning rules: the Hebbian and the BCM rule. Because of their spatially correlated activity, we proposed that peripherally generated L-events can establish topographically-refined receptive fields in the cortex, while H-events can regulate connection strength homeostatically. Although L-events successfully instructed topographic receptive field refinements in the Hebbian rule, naively including H-events provided too much depression, eliminating selectivity in the network despite fine-tuning. In contrast, in the BCM rule, H-events were indeed homeostatic, regulating the threshold between depression and potentiation. However, small L-events, which carry precise information for topographic connectivity refinements, mostly caused long-term depression in the synaptic weights disrupting topography. Inspired by the sliding threshold in the BCM rule, we proposed a similar adaptive mechanism operating at the single-cell level in the Hebbian rule, which regulates the amplitude of the cortically generated H-events according to the preceding average activity in the network to refine receptive fields with excellent topography. This mechanism adjusts the strength of H-events to homeostatically balance local increases and decrease in activity. Without any additional fine-tuning, this mechanism can also explain the long-term event sparsification of cortical activity as the circuit matures and starts responding to visual input. Therefore, we propose that L- and adaptive H-events cooperate to synergistically guide circuit organization of thalamocortical synapses during postnatal development.

### The origin of cortical event amplitude adaptation

After a re-examination of spontaneous activity recorded in the developing cortex *in vivo* between postnatal days 8 and 10 [26], we found evidence for the proposed H-event amplitude adaptation. This mechanism is sufficiently general in its formulation that it could be realized at the cellular, synaptic or network level. At the cellular level, the adaptation mechanism resembles intrinsic plasticity. Typically, intrinsic plasticity has been reported in response to long-term perturbations in activity or persistent changes in synaptic plasticity like LTP and LTD, where the intrinsic properties of single neurons are adjusted in an activity-dependent manner [48, 47]. During intrinsic plasticity, neurons can alter the number and expression levels of ion channels to adjust their input-output function either by modifying their firing thresholds or response gains, which could represent the substrate for H-event amplitude regulation. Our adaptation mechanism is consistent with fast intrinsic plasticity operating on the timescale of several spontaneous events supported by many experimental studies. For instance, intrinsic excitability of spine motoneurons is depressed after brief but sustained changes in spinal cord network activity in neonatal mice [50]. Similarly, hippocampal pyramidal neurons also exhibit a rapid reduction of intrinsic excitability in response to sustained depolarizations lasting up to several minutes [62]. In addition to reduced excitability, in the developing auditory system, enhanced intrinsic excitability has been reported in the cochlea followed reduced synaptic excitatory input from hair cells in a model of deafness, although this change is slower than our proposed adaptation mechanism [63].

At the synaptic level, our adaptation mechanism can be implemented by synaptic scaling, a process whereby neurons regulate their activity by scaling incoming synaptic strengths in response to perturbations [64]. A second possibility is short-term depression, which appears to underlie the generation of spontaneous activity episodes in the chick developing spinal cord [65, 66]. Similarly, release probability suppression has been reported to strongly contribute to synaptic depression during weak activity at the calyx of Held [67], which is more pronounced at immature synapses where morphological development renders synaptic transmission less effective [68, 69]. This is also the case in the cortex, where short-term synaptic plasticity in young animals is stronger [70]. Beyond chemical synapses, plasticity of gap junctions, which are particularly prevalent in development [71], could also be a contributing mechanism that adapts overall network activity [72, 73].

Finally, at the network level, the development of inhibition could be a substrate for amplitude adaptation of cortically generated events. The main inhibitory neurotransmitter, GABA, is thought to act as a depolarizing neurotransmitter, excitatory in early postnatal days [74], although recent evidence argues that GABAergic neurons have an inhibitory effect on the cortical network already in the second postnatal week [75, 76, 77]. Thus, the local maturation of inhibitory neurons – of which there are several types [78] – that gradually evolve to balance excitation and achieve E/I balance [79] could provide an alternative implementation of the proposed H-event adaptation.

### Developmental sparsification of cortical activity

On a longer timescale than receptive field refinement, we demonstrated that the adaptation of H-event amplitude can also bring about the event sparsification of cortical activity, as global, cortically generated H-events are attenuated and become more localized. The notion of ‘sparse neural activity’ has received significant attention in experimental and theoretical studies of sensory processing in the cortex, including differing definitions and implementations [80, 81, 59, 82, 83]. In particular, sparse activity in the mature cortex has been argued to be important for the efficient coding of sensory inputs of different modalities [82, 80]. Hence, the developmental process of receptive field refinement might be expected to produce sparser network activity over time. However, experiments directly testing this idea have found no, or even opposite, evidence for the developmental emergence of efficient sparse coding [59, 60]. In the context of our work, our notion of sparsification is fairly simplistic: it refers to an overall sparsification of network events (fewer active cells per event) where cortically-generated H-events become replaced with peripherally-driven L-events, following [26]. Given that our data pertain to developmental spontaneous activity before eye-opening, in complete absence of stimulation, it is not straightforward to relate our event sparsification to the sparse efficient coding hypothesis.

### Assumptions in the model

Our model was based on the assumption that L- and H-events have distinct roles during the development of the visual system. Retinal waves, the source of L-events, carry information downstream about the position and function of individual retinal ganglion cells [55], hence they are ideally suited to serve as ‘training patterns’ to enable activity-dependent refinements based on spatiotemporal correlations [84, 2, 4]. Since all cells are maximally active during H-events, these patterns likely do not carry much information that can be used for activity-dependent refinement of connectivity. In contrast, we assumed that H-events homeostatically control synaptic weights, operating in parallel to network refinements by L-events. Indeed, highly correlated network activity can cause homeostatic down-regulation of synaptic weights via a process known as synaptic scaling [39]. The homeostatic role of H-events is also consistent with synaptic downscaling driven by slow waves during sleep, a specific form of synchronous network activity [40, 85]. Since during development sleep patterns are not yet regular, we reasoned that refinement (by L-events) and homeostasis (by H-events) occur simultaneously instead of being separated into wake and sleep states.

We focused on the role of spontaneous activity in driving receptive field refinements, rather than study how spontaneous activity is generated. While the statistical properties of spontaneous activity in the developing cortex are well-characterized, the cellular and network mechanisms generating this activity remain elusive. In particular, while H-event generation has been shown to rely on gap junctions [26, 71], which recurrently connect developing cortical cells, not much is known about how the size of cortical events is modulated and how an L-event is prevented from spreading and turning into an H-event. It is likely that cortical inhibition plays a critical role in localizing cortical activity and shaping receptive field refinements [86, 35], for instance, through the plasticity of inhibitory connections by regulating E/I balance [79]. As new experiments are revealing more information about the cellular and synaptic mechanisms that generate spatiotemporally patterned spontaneous activity [87], a full model of the generation and the effect of spontaneous activity might soon be feasible.

We found that two learning rules can generate selectivity and receptive field refinement, the Hebbian rule with adaptive H-events, and the BCM rule. Both rules have an adaptive component: in the BCM rule it is the threshold between potentiation and depression that slides as a function of postsynaptic activity, while in the Hebbian rule it is the adaptive amplitude of H-events, while the rule itself is fixed. Although experimental data contains signatures of the stereotypical activity dependence of the BCM rule [88, 89], whether a sliding threshold exists is still debated. Moreover, the timescale over which the threshold slides to prevent unbounded synaptic growth needs to be much faster than experimentally found [90]. Our proposed H-event amplitude adaptation operates on the fast timescale of several spontaneous events found experimentally [26, 62, 50]. Hence, together with the better topography and the resulting event sparsification as a function of developmental stage that the Hebbian rule with adaptive H-events generates, we propose it as the more likely plasticity mechanism to refine receptive fields in the developing visual cortex.

Finally, we have focused here on the traditional view that molecular gradients set up a coarse map that activity-dependent mechanisms then refine [14]. In our model, this was implemented as a weak bias in the initial connectivity, which did not affect our results regarding the refinement of receptive fields. Both activity and molecular gradients may work together in interesting ways to refine receptive fields [91], and future work should include both aspects.

### Predictions of the model

Our model makes several experimentally testable predictions. First, we showed that changing the frequency of H-events can affect the size of the resulting receptive fields under both the BCM (Fig. 3) and the Hebbian rule with adaptive H-events (Fig. 4). The frequency of H-events can be experimentally manipulated using optogenetics or pharmacology. For instance, gap junction blocker (carbenoxolone) has been shown to specifically reduce the frequency of H-events [26], hence in that scenario our results predict broader receptive fields.

Additionally, L-events can also be experimentally manipulated. Recently, reduced inhibitory signaling by suppressing somatostatin-positive interneurons have has been shown to increase the size of L-events in the developing visual cortex [35]. With the effect of altered inhibitory signaling on receptive field refinements still unknown, our work predicts larger receptive fields and worse topography upon reduction of inhibition. L-events can also be experimentally manipulated by changing the properties of retinal waves, which can significantly affect retinotopic map refinement of downstream targets [52, 53, 12]. Indeed, *β*2 knockout mice discussed earlier have larger retinal waves and less refined receptive fields in the visual cortex [54, 55, 56]. If we assume that these larger retinal waves manifest as larger L-events in the visual cortex following [26], then these experimental observations are in agreement with our model results.

Third, our model predicts that as a result of receptive field refinement during development, network events sparsify as global, cortically generated events are attenuated and become more localized. Interestingly, the properties of spontaneous activity measured experimentally in different sensory cortices [33, 57, 92, 93, 94, 34] and in the olfactory bulb [87] change following a very similar timeline during development as predicted in our model. However, in many of these studies activity has not been segregated into peripherally driven L-events and cortically generated H-events. Therefore, our model predicts that the frequency of L-events would increase while the frequency of H-events would decrease over development.

Finally, we propose that for a Hebbian rule to drive developmental refinements of receptive fields using spontaneous activity L- and H-event patterns recorded *in vivo* [26], H-events need to adapt to ongoing network activity. Whether a fast adaptation mechanism like the one we propose operates in the cortex requires prolonged and detailed activity recordings *in vivo*, which are within reach of modern technology [2, 95, 25]. Our framework also predicts that manipulations that affect overall activity levels of the network, such as activity reduction by eye enucleation, would correspondingly affect the amplitude of ongoing H-events.

### Conclusion

In summary, we studied the refinement of receptive fields in a developing cortex network model constrained by realistic patterns of experimentally recorded spontaneous activity. We proposed that adaptation of the amplitude of cortically generated spontaneous events achieves this refinement without additional assumptions on the type of plasticity in the network. Our model further predicts how cortical networks could transition from supporting highly-synchronous activity modules in early development, to sparser peripherally driven activity suppressing local amplification which could be useful for preventing hyper-excitability and epilepsy in adulthood, while enhancing the processing of sensory stimuli.

## Methods and Materials

### Network model

We studied a feedforward, rate-based network with two one-dimensional layers, one of *N*_*u*_ thalamic neurons (u) and the other of *N*_*v*_ cortical neurons (**v**), with periodic boundary conditions in each layer to avoid edge effects. The initial connectivity in the network was all-to-all with uniformly distributed weights in the range *w*_ini_ = [*a, b*]. In addition, a topographic bias was introduced by modifying the initially random connectivity matrix to have strongest connections between neurons at the matched topographic location, and which decay with a Gaussian profile with increasing distance (Fig. 2C), with amplitude *s* and spread *σ*_*s*_. During evolution of the weights, soft bounds were applied on the interval [0, *w*_max_]. We studied weight evolution under two activity-dependent learning rules: the Hebbian (Eq. 2) and the BCM (Eq. 3) rules. Table 1 lists all parameters. Sample codes can be found at github.com/comp-neural-circuits/LH-events.

### Generation of L- and H-events

We modeled two types of spontaneous events in the thalamic (L-events) and the cortical (H-events) layer of our model [26]. During L-events, the firing rates of a fraction (*L*_pct_) of neighboring thalamic neurons were set to *L*_amp_ = 1 during a period *L*_dur_ and were otherwise 0. Similarly, during H-events, the firing rates of a fraction (*H*_pct_) of cortical cells were set to *H*_amp_ during time *H*_dur_. As a result, cortical neuron activity was composed of H-events and L-events transmitted from the thalamus. For each H-event, *H*_amp_ was independently sampled from a Gaussian distribution with mean *H*_amp_ and standard deviation *H*_amp_*/*3. The inter-event intervals were *L*_int_ and *H*_int_ sampled from experimentally characterized distributions in [26] (Table 1). We note that in the experiments, both L- and H-events were characterized in the primary visual cortex; in our model, we assume that L-events are generated in the retina and subsequently propagated through the thalamus to the cortex, where they manifest with the experimentally reported characteristics (see Fig. 7B for example). This interpretation is supported by experimental evidence [26], but we cannot exclude the possibility that the retina also generates H-events, or that L-events are generated in the cortex.

### Reduction of the weight dynamics to two dimensions

To reduce to reduce the full weight dynamics to a two-dimensional system, we averaged all the *n* weights belonging to the receptive field that are predicted to potentiate along the initial topological bias, as *w*_RF_, and all the *N*_*u*_ − *n* remaining weights, which we call complementary to the receptive field, as *w*_C_. When all weights behaved the same, we arbitrarily split them into two groups of the same size. Details about the classification of weights as *w*_RF_ or *w*_C_ can be found in the Appendix.

### Computing the strength of simulated H-events

To relate the reduced two-dimensional phase planes to the simulation results, we wrote down the steady state activity of neuron *j* Eq. 1, which contains the rate gain from H-events relative to L-events, ⟨*R*_H_⟩ (also called ‘Strength of H-events’ in Fig. 4E) :

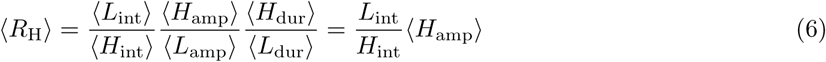

since ⟨*L*_dur_⟩ = ⟨*H*_dur_⟩ and ⟨*L*_amp_⟩ = 1.

In the absence of adaptive H-events, for a fixed set of values for *H*_amp_ and *L*_int_ (as in Table 1) and a chosen ⟨*R*_H_⟩ which we called ‘Non-adapted strength of H-events’ in Fig. 4E, we used Eq. 6 to find the *H*_int_ value that satisfies the equation. Next, we ran simulations with the same *H*_int_ and *L*_int_ parameters, but adaptive *H*_amp_. We fixed the inter-event intervals of both L- and H-events to their mean values *L*_int_ and *H*_int_ instead of sampling them from distributions in Fig. 4E. Then we numerically estimated the average amplitude of H-events with adaptation which we called ‘Adapted strength of H-events’ in Fig. 4E at the end of the simulation (final 5% of the simulation time) when the dynamics were stationary.

### Receptive field statistics

The following receptive field statistics were used to quantify properties of the weight matrix ***W*** after the developing weights became stable.

#### Receptive field size

The receptive field of a cortical neuron is the group of weights from thalamic cells for which *w*_*ij*_ *> w*_max_*/*5. The lower threshold was chosen to make the measurement robust to small fluctuations around 0, which are present because of the soft bounds. Mathematically, we compute the receptive field size of cortical neuron *j* as:

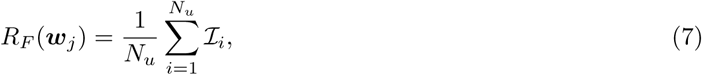

with the ***ℐ*** vector given by:

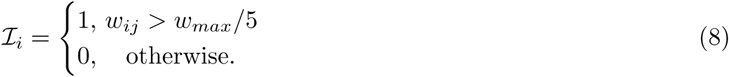

The normalized receptive field ranges from 0 corresponding to a total decoupling of the cortical cell from the input layer, to 1 corresponding to no selectivity due to the potentiation of all weights from the input layer to that neuron. To compute the average receptive field size of the network, we include only the cortical neurons (*N* ^*\**^) that have not decoupled:

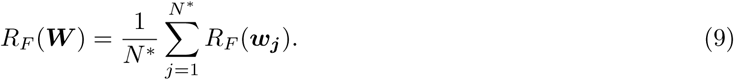

If all the cortical cells have decoupled from the thalamus, we set *R*_*F*_ (***W***) = 0.

#### Topography *𝒯*

The topography of the network is a measure of how much of the initially weak biased topography is preserved in the final receptive field. Due to our biased initial conditions, neighboring thalamic cells are expected to project to neighboring cortical cells, yielding a diagonal weight matrix. For each cortical neuron, we calculated how far the center of its receptive field is from the ideal diagonal. Mathematically, for each row *j* of ***W***, we determined the center of the receptive field *c*_*j*_ and calculated the smallest distance (while considering periodic boundary conditions) between the receptive field center and the diagonal element *j*. Then, we summed all the squared distances and calculated the average error of the topography:

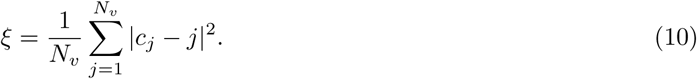

To normalize the topography, we compared *ξ* to the topography error Ξ of a column receptive field (Fig. S1A) where the centers of all cortical receptive fields were the same, *c*_*j*_ = *c* (*c* a constant). For such a column receptive field, 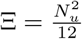. Therefore, we define the topography score *𝒯* as:

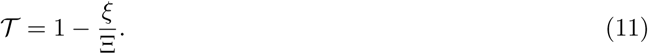

The topography will be close to 1 if the weight matrix is perfectly diagonal, and 0 if the final receptive field is a column (*ξ* = Ξ).

#### Proportion of cortical decoupling *D*

To quantify the cortical decoupling we use Eq. 8 to compute the fraction of decoupled neurons divided by the number of neurons, 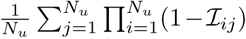. If the decoupling is 0, no cortical neuron has decoupled from the thalamus, while decoupling of 1 means that all the cortical neurons are decoupled from the thalamus.

### Quantifying adaptation in the data

We first investigated if fluctuations in the activity across recordings could generate significant correlations. We analyzed consecutive recordings (each *∼*5 mins long) in the same animal of which we had between 3 and 14 in all 26 animals (separated by ¡ 5 mins due to experimental constraints on data collection) to identify possible fluctuations on a longer timescale. We found that the average amplitude of all (L and H) events is not significantly different across consecutive recordings of the same animal (Fig. S3A, one-way ANOVA tests, *p >* 0.05 in 23 out of 26 animals). Across different animals and ages, individual event amplitudes remained uncorrelated between successive recordings at this timescale, which we confirmed by plotting the difference in event amplitude as a function of the time between recordings (Fig. S3B), Kruskal-Wallis test, *p >* 0.05.

For our reanalysis of the spontaneous events (Fig. 5), we only included events that recruited at least 20% of the cells in the imaging field of view following [26]. We computed the average amplitude of all events that occurred within a time window *τ*_max_ before an H-event (consecutive recordings were concatenated) and compared it to the amplitude of the H-event. We excluded animals that had fewer than 12 H-events preceded by spontaneous activity within the time window *τ*_max_ (9 animals remained after exclusion). Next, we computed the correlation coefficient of the relationship between H-event amplitude and the average amplitude of preceding activity within *τ*_max_ with a leaky accumulator of time constant *τ*_decay_. To estimate the 95% confidence interval, we performed a bootstrap analysis in which we generated 1000 bootstrap datasets by drawing without replacement from the valid pairs of H-event amplitudes and average amplitude of preceding activity. We repeated this analysis with different thresholds for excluding data (Fig. S4A,B), different values of the time window *τ*_max_ within which events are averaged (Fig. S4C) and for different decay time constants *τ*_decay_ (Fig. S4D). All data and analysis code can be found at github.com/comp-neural-circuits/LH-events.

### Spontaneous events identification in the model

To quantify the properties of spontaneous activity in the cortical layer of our model, we used the time series of activity of all the simulated cortical neurons (after weight stabilization is achieved) sampled in a high time resolution (0.01 s, Fig. 7B). We defined a global activity threshold *ν* = *v*_max_*/r*, where *v*_max_ is the highest amplitude among the cortical cells in the recording and *r* is a fixed scaling constant (*r* = 8 for all recordings). For each cortical cell *j*, we labeled the intervals where the cell was active (1) or inactive (0) based on:

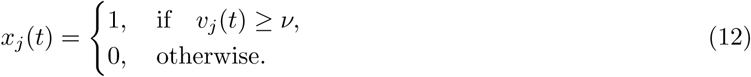

We then used the trace 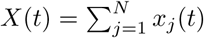 to define the number of active cortical cells at each time step *t*. If at time step *t* = *τ* the proportion of active cells is between 20-80% of the total number *N*_*v*_, then we label *X*(*τ*) as an effective L-event, otherwise if the proportion of active cells is higher than 80%, we label *X*(*τ*) as an effective H-event. The duration of each event was then the number of continuous time steps classified as either L- or H-event, and the inter-event interval was calculated based on the number of time steps between two events of the same type. For each identified event, we averaged the amplitude of the active cells to obtain the amplitude vs. participation rate relationship.

## Acknowledgments

This project has received funding from the European Research Council (ERC) under the European Union’s Horizon 2020 research and innovation programme (Grant agreement No. 804824 to JG). This work was further supported by the Max Planck Society (MW, JHK, LYC, JG), a NARSAD Young Investigator Grant from the Brain and Behavior Research Foundation (JG), a Capes-Humboldt Research fellowship (MW), the Smart Start joint training program in computational neuroscience (JHK), and by grants of the Netherlands Organization for Scientific Research (NWO, ALW Open Program grants, no. 819.02.017, 822.02.006 and ALWOP.216; ALW Vici, no. 865.12.001) and the “Stichting Vrienden van het Herseninstituut” (NZ, CL). We thank Stephen Eglen for discussions and ideas in the initial stages of the project.

## Appendix

### The weight dynamics under the Hebbian rule

Since synaptic plasticity operates on a much slower time scale than the response dynamics of the output neuron, we make a steady state assumption and write Eq. 1 for neuron *j* as:

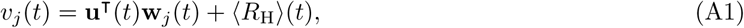

where ***w***_*j*_ is the vector of the elements in the row of the weight matrix ***W***, i.e. the vector of weights from all thalamic neurons into cortical neuron *j*. ⟨*R*_H_⟩ (*t*) is the rate gain from H-events relative to L-events, which depends on the duration, amplitude, and inter-event intervals of both L- and H-events. Specifically, ⟨*R*_H_⟩ is proportional to ⟨*H*_amp_⟩, ⟨*H*_dur_⟩, ⟨*L*_int_⟩ and inversely proportional to ⟨*H*_int_⟩, ⟨*L*_amp_⟩ and ⟨*L*_dur_ ⟩ (where ⟨ *·* ⟩denotes an ensemble average over the activity patterns), such that:

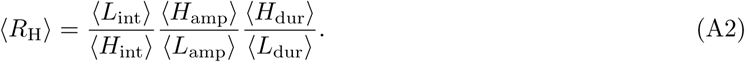

Rewriting Eq. 2 in vector form:

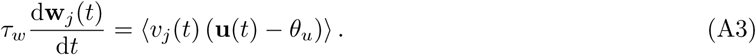

Inserting Eq. A1 into the weight dynamics from Eq. A3 yields (vector dependence on *t* is omitted for clarity):

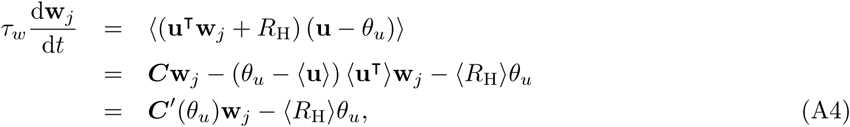

where ***C*** = ***Q*** − ⟨*u*⟩2 is the input covariance matrix, ***Q*** = ⟨***uu*^T^**⟩ is the input correlation matrix and we have defined ***C*′**(*θ*_*u*_) = ***C*** − ⟨*u*⟩ (*θ*_*u*_ − ⟨*u*⟩) = ***Q*** − ⟨*u*⟩*θ*_*u*_ to be the ‘modified covariance matrix’ of the learning rule. We write ⟨***u***⟩ to denote the vector with repeated element ⟨*u*⟩, which is the mean normalized size of an L-event (e.g., for L-events engaging 20–80% of input neurons, ⟨*u*⟩ = 0.5). We also used the fact that the occurrence of L- and H-events is uncorrelated [26], such that ⟨*R*_H_***u***⟩ = 0.

### The input correlation matrix

The spontaneous L-events in our network model have useful mathematical properties that allowed us to derive an analytical expression for the elements of the input correlation matrix ***Q***. The locality and periodicity of L-events, as well as the long-term averaging of their stationary dynamics generate a symmetric and circulant matrix ***Q*** [96]. In a circulant matrix, each row is rotated one element to the right relative to the preceding row, and thus ***Q*** can be completely defined by a vector.

Let *ℒ* be the fixed size of an L-event between 0 and *N*_*u*_. By computing the correlations among all the possible L-events of size *ℒ*, we can write the elements of the vector 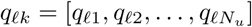, which completely defines ***Q***, as:

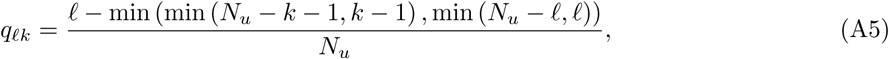

where min(*a, b*) returns the minimum of *a* and *b*. Since we wanted to explore the dependence of refinement on the size of L-events, defined by the minimum (*ℒ*_min_) and maximum (*ℒ*_max_) number of cells they activate in the input layer, we average over the size of a given event:

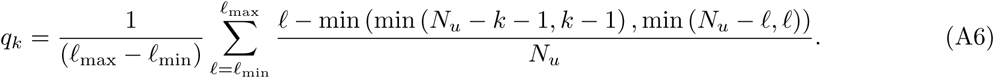

Finally, using the emergent symmetry of ***Q***, we can simplify the elements of this vector as follows:

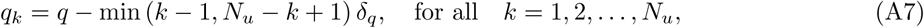

with q = (*l*_*max*_ + *l*_*min*_)/2*N*_*u*_ and *δ*_*q*_ = 1/*N*_*u*_

Since ***C***(*θ*_*u*_) and ***C*′**(*θ*_*u*_) are defined in terms of ***Q*** minus a constant, both are circulant matrices as well.

### Fixed point of the weight dynamics

The fixed point of the weight dynamics under L- and H-events is obtained by setting Eq. A4 to zero and solving the resulting equation for ***w***^***\****^:

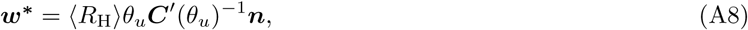

where ***n*** is a vector of ones. To study the nature of the fixed point ***w***^***\****^, we need to investigate the eigenvalues of the circulant matrix ***C*′**(*θ*_*u*_)^−1^ (the inverse of a circulant matrix is also a circulant matrix). A circulant matrix has the property that its eigenvectors can be written in terms of roots of unity. The eigenvalues are real and can be written as the discrete Fourier transforms of any row of the matrix [96]. In particular, one of the eigenvalues is the sum of the elements of any row of the matrix, with a corresponding eigenvector that is a constant. We call this special eigenvalue the ‘row-sum eigenvalue’. All eigenvalues except for the row-sum are non-negative.

We can write the fixed point as ***w***^***\****^ = (*λ*^*\**^)^−1^*θ*_*u*_ ⟨*R*_H_⟩ ***n***, where *λ*^*\**^ is the row-sum eigenvalue of ***C*′**(*θ*_*u*_). It is clear that if only L-events are present, ⟨*R*_H_⟩ = 0, the fixed point is always at the origin, ***w***^***\****^ = 0. Consequently, adding H-events in the cortical layer moves the location of the fixed point along the diagonal of the phase plane, i.e. equally in all directions. The fixed point will be positive (negative) when *λ*^*\**^ is positive (negative). Furthermore, it will be an unstable fixed point for *λ*^*\**^ > 0, since all the eigenvalues of the dynamical system are positive. The fixed point is a saddle node when *λ*^*\**^ < 0 since the other eigenvalues are positive.

### Calculation of receptive field size

The Hebbian rule on its own does not have a mechanism to prevent the weights to grow infinitely large or negative. Thus, to generate receptive fields we imposed a lower bound at 0 and an upper bound at *w*_max_. This enabled us to calculate the size of the receptive field, *n*, as a function of L-event properties and the input threshold, *θ*_*u*_ in the absence of H-events. Because the cortical cells are not recurrently connected, here we examine the weight vector onto a single cortical neuron. Assuming the dynamical system has reached steady state, to get a receptive field of size *n < N*_*u*_ for this cortical neuron, we can write the fixed point weight vector as ***w***^***\****^ = (*w*_max_, *…, w*_max_, 0, *…*, 0)^T^, where *n* of the weights have reached the upper bound *w*_max_, while the remaining weights the lower bound 0. The specific weight identity reaching the upper bound is not important, as long as they are topographically near each other. To achieve this fixed point:

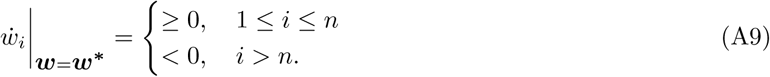

Using the structure of our circulant correlation matrix (Eq. A7), we can rewrite Eq. A4 (note that in the absence of H-events ⟨*R*_H_⟩ = 0) as:

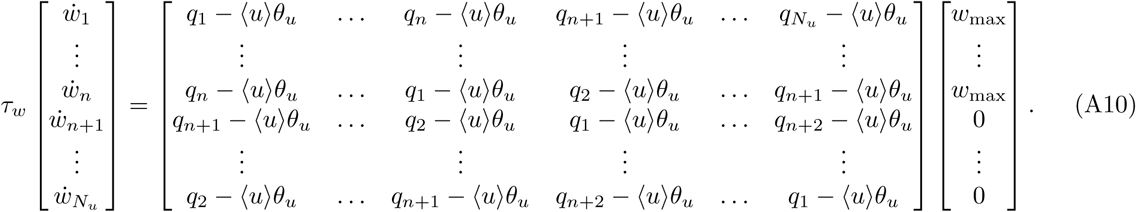

Now, we study the conditions to guarantee that Eq. A9 is satisfied. Computing each equation explicitly, we obtain:

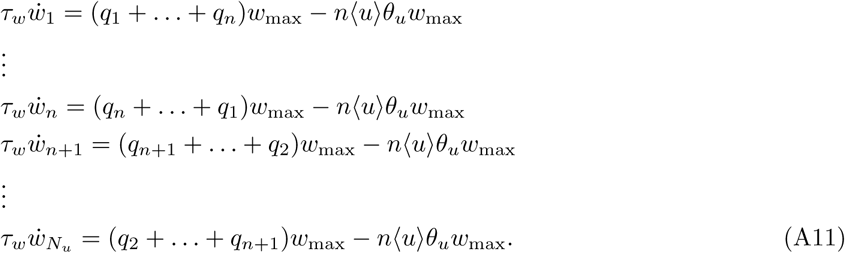

And finally, using Eq. A7 and after some manipulations this can be written as:

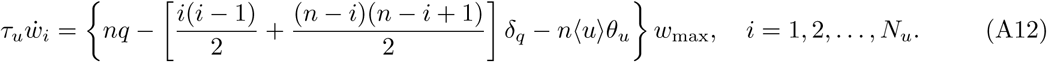

To determine the input threshold *θ*_*u*_ in Eq. A12 that yields a receptive field of size *n*, we assume that 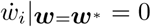 for *i* = *n*; this implies that 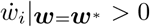 for *i* < n and 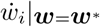 < 0 for *i > n*. We write *θ*_*u*_ as a linear combination of *q* and *δ*_*q*_:

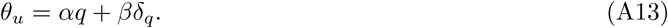

Using this ansatz in Eq. A12 and setting it to zero for *i* = *n* (since *w*_max_ *>* 0), we find that *α* = 1*/*⟨ *u*⟩ and *β* = − (*n* − 1)*/*(2 ⟨*u*⟩). Therefore, only in the presence of L-events, we derive the input threshold at which the resulting receptive field size is *n*:

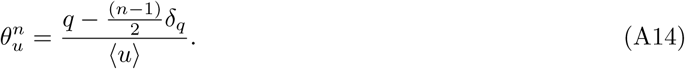

Plugging this threshold into Eq. A12 for all *i*, we get a quadratic polynomial in *i*:

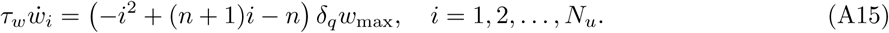

Since *δ*_*q*_, *w*_max_ *>* 0, indeed 1 ≤ *i* ≤ *n* results in 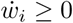 while *i > n* yields 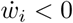, thus, satisfying Eq. A9.

In the absence of H-events, we computed the average size of receptive fields using Eq. A14 for a range of input thresholds *θ*_*u*_ and maximum participation rates of L-events, while keeping the minimum participation rate at 20% (Fig. A1A, contour lines). We verified our analytical predictions with Monte Carlo simulations for the same range of parameters (Fig. A1A). We confirmed that the size of L-events, which depends on the range of participation rates, has a direct impact on receptive field size, with larger L-events resulting in larger receptive fields for a fixed input threshold (Fig. A1B). Low input thresholds generate refined receptive fields only if the size of spontaneous events is small.

### The eigenspace of *C*′(*θ*_*u*_)

To gain intuition for the weight dynamics in Eq. A4, we first investigated the eigenspace of ***C*′**(*θ*_*u*_), the vector space spanned by the eigenvectors of ***C*′**(*θ*_*u*_). Specifically, we focused on the conditions that enabled the robust formation of cortical receptive fields.

Using the fact that ***Q*** and ***C*′**(*θ*_*u*_) are circulant matrices (Fig. A2A, B), we identified two input thresholds, *θ*^*\**^ and *θ*^*\*\**^, that define three dynamical regions that the row-sum eigenvalue of ***C*′**(*θ*_*u*_), *λ*^*\**^, can occupy (Fig. A2C). To obtain the first critical input threshold *θ*^*\**^ that characterizes the transition from region (i) to region (ii), we set, for any row *j*, the row-sum eigenvalue to the largest (fixed) eigenvalue of ***C*′**(*θ*_*u*_):

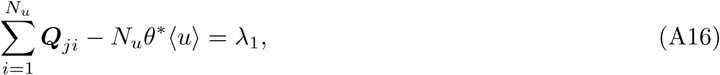

and for the input statistics of experimentally measured L-events (Table 1), we obtained *θ*^*\**^ = 0.414. Similarly, the transition from region (ii) to region (iii) is achieved when the row-sum eigenvalue is set to zero:

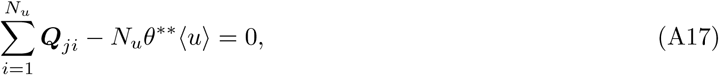

and the second critical input threshold is obtained as *θ*^*\*\**^ = 0.564.

**Figure A1.**
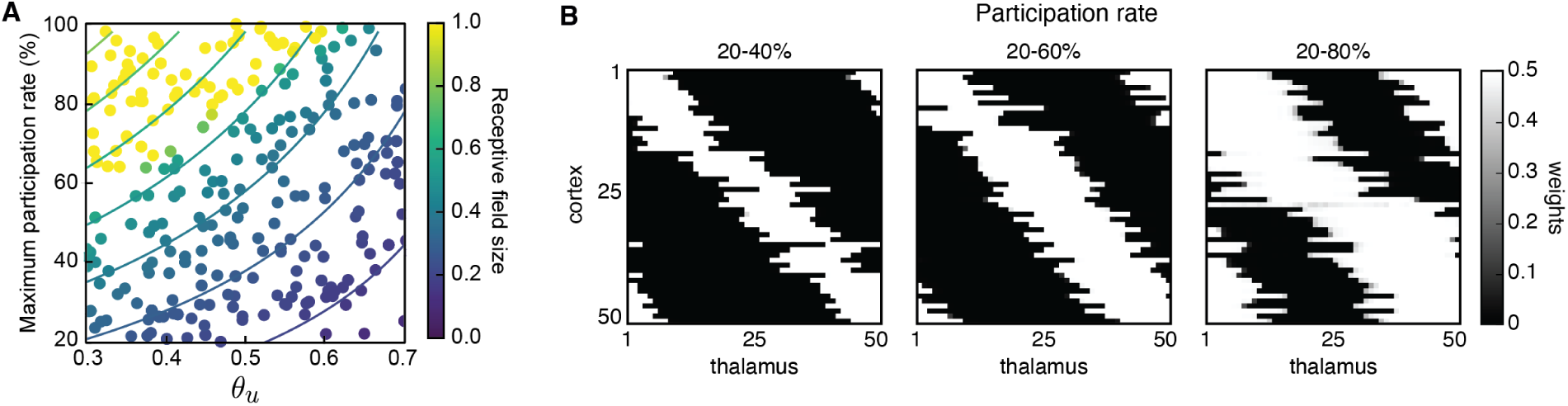
Receptive field size depends on L-event properties and learning rule input threshold in the absence of H-events. **A**. Receptive field sizes obtained from 500 Monte Carlo simulations for combinations of L-event maximum participation rate and input threshold, *θ*_*u*_. For all simulations, the L-event minimum participation rate was fixed at 20%. The contour plots of receptive field sizes were obtained using the analytical approach (Eq. A9). B. Example receptive fields for different L-event maximum participation rates and *θ*_*u*_ = 0.5. Smaller events recruiting only 20–40% of the input neurons generate very refined receptive fields. As the upper bound of the participation rate progressively increases from 40% to 80%, receptive fields get larger.

In region (i), 0 < *θ*_*u*_ < *θ*^*\**^ and *λ*^*\**^ *>* 0 is the dominant (largest) eigenvalue of ***C*** (*θ*_*u*_). The eigenvector corresponding to it is a constant (Fig. A2D, (i)). This predicts that all synaptic weights to a given cortical cell will potentiate preventing the formation of a localized receptive field. Since all eigenvalues are non-negative, the fixed point is an unstable node of the linear dynamical system (Eq. A4).

In region (ii), *θ*^*\**^ *< θ*_*u*_ < *θ*^*\*\**^ and *λ*^*\**^ *>* 0. However, *λ*^*\**^ is no longer the dominant eigenvalue. In this setting, there is a pair of dominant eigenvalues with the corresponding eigenvectors taking the form of out-of-sync sine waves with positive and negative elements. The sign of these elements predicts that some weights will potentiate while others depress, thus enabling the formation of receptive fields (Fig. A2D, (ii)). All eigenvalues in this second region remain non-negative and the fixed point is still an unstable node of the dynamical system.

Finally, in region (iii), *θ*^*\*\**^ < *θ*_*u*_ < 1 and *λ*^*\**^ < 0. While the dominant eigenvectors are similar to those in region (ii), enabling the formation of localized receptive fields (Fig. A2D, (iii)), the dynamics of the dynamical system are different because the fixed point is now a saddle node.

### Analysis of the weight dynamics in two dimensions

We reduced the dimension of the weight dynamics by defining two distinct sets of weights. The first set, *w*_RF_, corresponds to the *n* weights which correspond to the topographically biased locations of the receptive field. The complementary set *w*_C_ contains the remaining weights. To classify the weights into *w*_RF_ and *w*_C_, first we solved Eq. A4 with the biased initial condition and limited the sum to the dominant eigenvalues and respective eigenvectors. In the case of selectivity, the eigenvectors have half the elements positive and half negative. Therefore, if the receptive field size is *n* < *N*_*u*_*/*2 we needed to subsample the potentiating weights to achieve the smaller receptive field size. We did it by keeping only the *n* largest positive elements in *w*_RF_ and moving the remaining ones to *w*_C_. If *n > N*_*u*_*/*2, we downsampled *w*_C_ by moving the *n* − *N*_*u*_*/*2 less-negative weights to *w*_RF_. Due to the topographically biased initial conditions, *w*_RF_ always contained the weights potentiating along the diagonal *w*_RF_ = *w*_C_.

We then regularly sampled initial conditions in [0, *w*_max_] *×* [0, *w*_max_]. For a given receptive field size *n*, we set 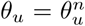 according to Eq. A14. If *w*_RF_(0) *> w*_C_(0), *w*_RF_ contains the *n* weights that form the receptive field by potentiating to the upper bound *w*_max_, while *w*_C_ contains the remaining *N*_*u*_ − *n* weights that depress to 0. Similarly, if *w*_RF_(0) < *w*_C_(0), *w*_C_ contains the *n* weights that potentiate to the upper bound, while *w*_RF_ contains the remaining *N*_*u*_ − *n* weights that depress to 0. At each initial condition, we solved the weight evolution (Eq. A4) for each weight, and averaged the weights in *w*_RF_ and *w*_C_ for a small time interval to obtain the direction of the phase plane arrows. We computed the evolution trajectory by solving Eq. A4 with a topographically biased initial condition.

**Figure A2.**
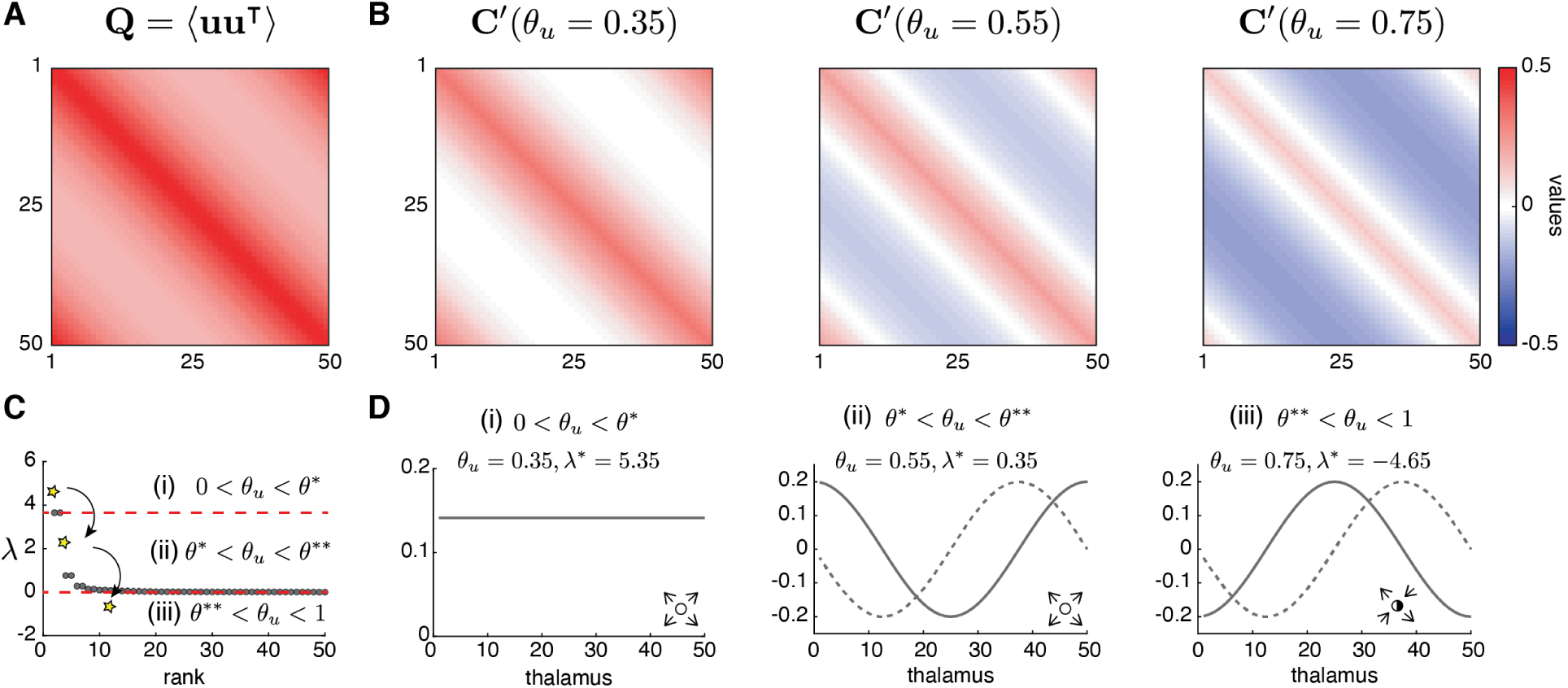
Eigenvalues and eigenvectors of weight dynamics predict receptive field refinement. **A**. The input correlation matrix ***Q*** = ⟨***uu***^**T**^⟩. **B**. The modified covariance matrices ***C*′**(*θ*_*u*_) = ***Q*** − ⟨*u*⟩ *θ*_*u*_ for different input thresholds, *θ*_*u*_. **C**. Two thresholds, *θ\**= 0.414 and *θ\*\** = 0.564, define three different dynamical regions in the spectrum of ***C*′**(*θ*_*u*_), delineated by the horizontal red dashed lines: (i) 0 < *θ*_*u*_ < *θ*^*\**^, (ii) *θ*^*\**^ < *θ*_*u*_ *< θ*^*\*\**^ and (iii) *θ*^*\*\**^ *< θ*_*u*_ *<* 1. The row-sum eigenvalue *λ*^*\**^ in each case is given by the yellow star, while the remaining eigenvalues are shown as gray circles. **D**. Dominant eigenvectors corresponding to each region in C. Inset: fixed points corresponding to each region: (i), (ii) unstable node (open circles); (iii) saddle node (half-open circle).

### Analytical solution of the two-dimensional weight dynamics with only L-events in the Hebbian rule

We first examined the weight dynamics in the reduced two-dimensional phase plane *w*_RF_ *× w*_C_ for only L-events (⟨*R*_H_⟩ = 0 in Eq. A4). The phase plane is symmetric about the diagonal *w*_RF_ = *w*_C_ due to the symmetry of the dominant eigenvectors (Fig. A3A). As predicted by the eigenvectors (Fig. A2D), in region (i) both *w*_RF_ and *w*_C_ converge to the upper bound and the fixed point, which is located in the origin, is an unstable node (Fig. A3A, left). Therefore, all weights potentiate and no receptive field can be formed. In regions (ii) and (iii), the eigenvectors predict the formation of receptive fields with *w*_RF_ *→ w*_max_ and *w*_C_ *→* 0, respectively (Fig. A3A, middle and right), though the dynamics are different in each case because the origin is an unstable node or a saddle node, respectively.

Our analytical predictions of the reduced two-dimensional system with only L-events were confirmed in numerical simulations of the full *N* -dimensional system. In particular, in region (i) all weights potentiate with each cortical cell receiving input from all thalamic inputs, such that no receptive field forms (Fig. A3B, left). In regions (ii) and (iii), receptive fields form with good topography (Fig. A3B, middle and right). Therefore, consistent with the analytical prediction of receptive field size (Fig. A1), higher input thresholds resulted in smaller receptive fields.

Thus, the Hebbian rule can generate receptive fields of size depending on the input threshold *θ*_*u*_ in the presence of only L-events originating from the sensory periphery. This result is in agreement with previous findings of the emergence of other aspects of development including topographic maps [61] and ocular dominance [29, 30] or other selectivity [41, 97, 31] in the presence of correlated activity in the input layer of similar feedforward networks. We find that when the only input to the cortex are peripheral L-events, intrinsic properties of the learning rule, such as the threshold between potentiation and depression, control receptive field refinement.

**Figure A3.**
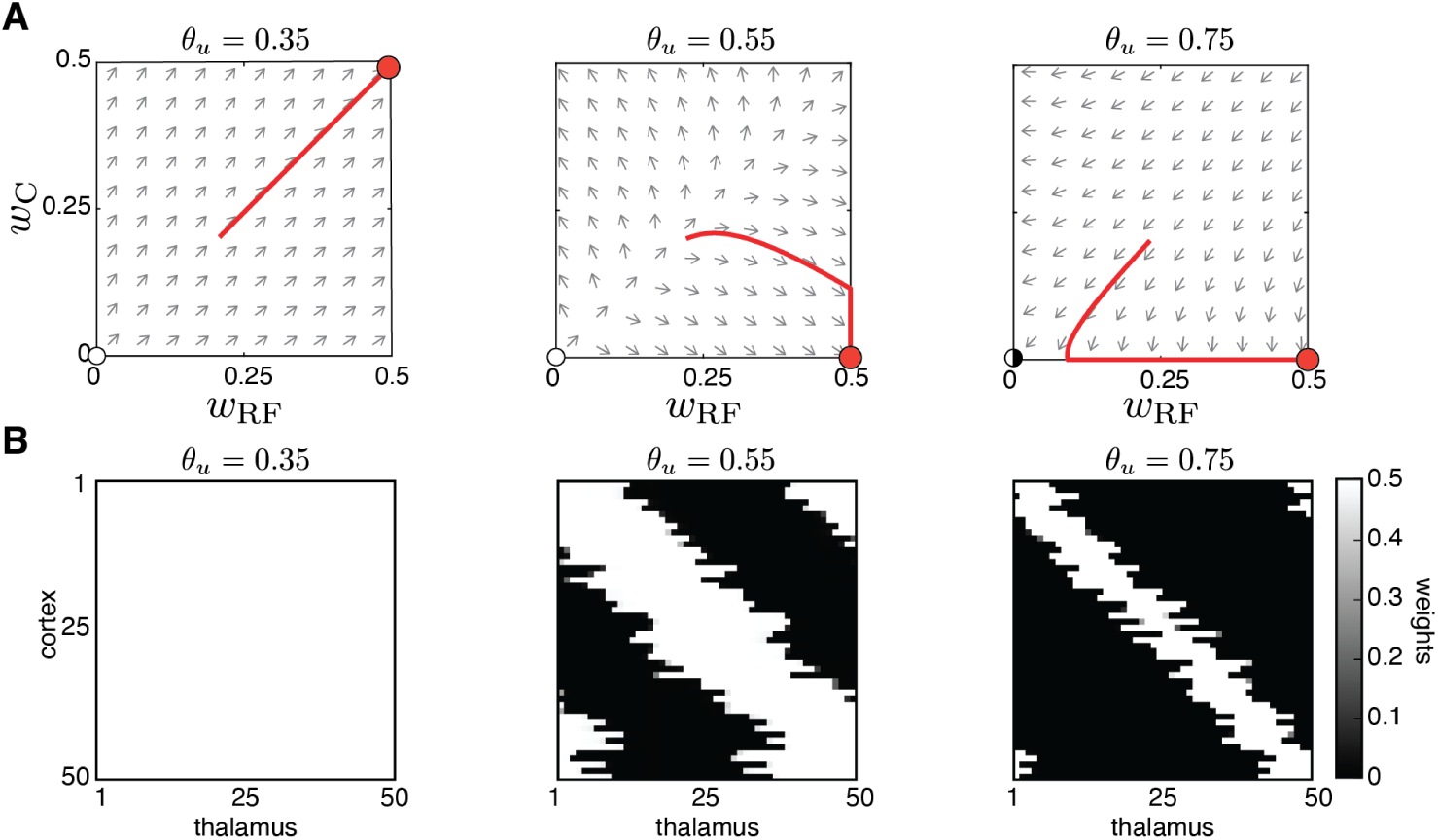
Peripheral L-events generate robust receptive field refinement. **A**. The reduced two-dimensional weight dynamics in the phase plane with the same *θ*_*u*_ as Fig. A2B and D. For each plane, the red trajectory depicts the weight evolution from an initial condition where *w*_RF_(0) *> w*_C_(0) until the weights’ upper bound (*w*_max_ = 0.5). Left: *w*_C_ *→ w*_max_ and *w*_RF_ *→ w*_max_, resulting in no selectivity. Middle and right: *w*_C_ *→* 0 and *w*_RF_ *→ w*_max_, resulting in selectivity and receptive field refinement. **B**. Simulation results for the same input thresholds of Fig. A2B, D for the full 50-dimensional system.

### Analytical solution of the two-dimensional weight dynamics with L- and H-events in the Hebbian rule

We also studied how the addition of H-events affects network refinements. To investigate the role of H-events in a systematic way, we repeated our analytical study of the weight-dynamics from Eq. A4, but with ⟨*R*_H_⟩≠ 0. In the reduced two-dimensional phase plane, *w*_RF_ ×*w*_C_, including spontaneous events in the cortical layer moves the fixed point of Eq. A4 away from the origin to the coordinates *w*_RF_ = *w*_C_ = (*λ*^*\**^)^−1^*θ*_*u*_ ⟨*R*_H_⟩. Nevertheless, the different dynamical regimes reported in Fig. A2C continue to be valid. In region (i), the addition of cortical events moves the unstable node away from the origin and into the first quadrant (Fig. A4A top). As a result, a small region of selectivity emerges in the plane in which initial conditions generate refined receptive fields (Fig. A4A, top middle). Therefore, the addition of H-events enables the emergence of weight selectivity but through a different mechanism than the one obtained with only L-events. Rather than modulating the learning rule through the input threshold, changing the H-event statistics through the ⟨*R*_H_⟩ parameter can generate different receptive field sizes for a fixed input threshold. However, the strength of H-events has to be fine-tuned to generate refined receptive fields. Within a small range of ⟨*R*_H_⟩ the network transitions from no-selectivity (where all weights potentiate, Fig. A4A, top left) to complete decoupling (where all weights depress, Fig. A4A, top right).

To relate the reduced two-dimensional phase planes to the simulation results, we used Eq. A2 to obtain ⟨*R*_H_⟩ by taking into account the simulation parameters in Table 1. We next verified the predictions of the reduced two-dimensional system in numerical simulations of the full network with H-events (Fig. A4A bottom). To capture the gradual increase of ⟨*R*_H_⟩ as in the reduced two-dimensional system, we decreased the average inter-event interval between H-events, *H*_int_. As before, only a narrow range of *H*_int_ leads to refined receptive fields, albeit with some degree of decoupling (Fig. A4A, bottom middle). Outside of this range, individual cortical neurons are either non-selective (Fig. A4A, bottom left) or nearly completely decoupled from the thalamus (Fig. A4A, bottom right). In regions (ii) and (iii), the fixed point moves from the origin to the first and third quadrants, respectively (Fig. A4B and Fig. A4C). In both cases, only very weak H-events can sustain finite receptive fields because the high input threshold value already provides sufficient depression to the network. We confirmed our analytical results in regions (ii) and (iii) with numerical simulations of the full network (Fig. A4B, bottom and Fig. A4C, bottom).

**Figure A4.**
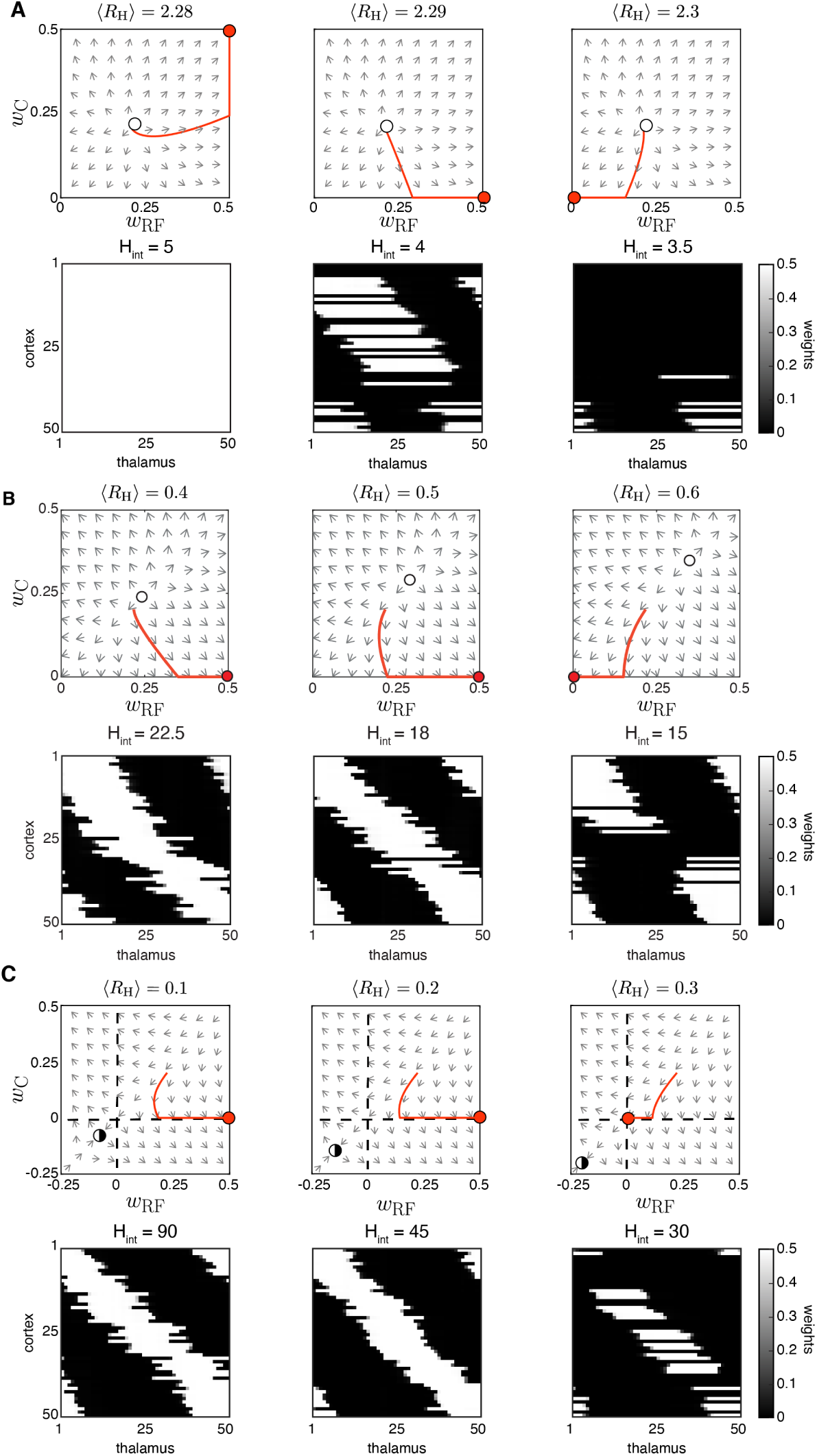
Spontaneous cortical H-events disrupt receptive field refinement. **A**. Top: Phase planes of the reduced two-dimensional system for input threshold *θ*_*u*_ = 0.4 (region i) and increasing strength of cortical events ⟨*R*_H_⟩ with an example trajectory (red). Selectivity can only be observed for fine-tuned ⟨*R*_H_⟩. The fixed point (open circle), an unstable node, has moved to the first quadrant. Bottom: Simulations of receptive field development with the same parameters where *H*_int_ was progressively reduced (this is the same set of parameters as shown in Fig. 3A). **B**. Top: Phase planes for *θ*_*u*_ = 0.52 (region ii) and increasing ⟨*R*_H_⟩ with an example trajectory (red). The unstable node (open circle) moves from the origin to the first quadrant as ⟨*R*_H_⟩ increases. Bottom. Simulations with the same parameters where *H*_int_ was progressively reduced. **C**. Top: Phase planes for *θ*_*u*_ = 0.6 (region iii) show the transition from selective receptive fields to cortical decoupling in response to increasing ⟨*R*_H_⟩. The fixed point (half-open circle), now a saddle node because *λ*^*\**^ *<* 0, has moved away from the origin to the third quadrant. Bottom: Simulations with the same parameters with very infrequent H-events where *H*_int_ was progressively reduced.

## Supplementary Figures

**Figure S1.**
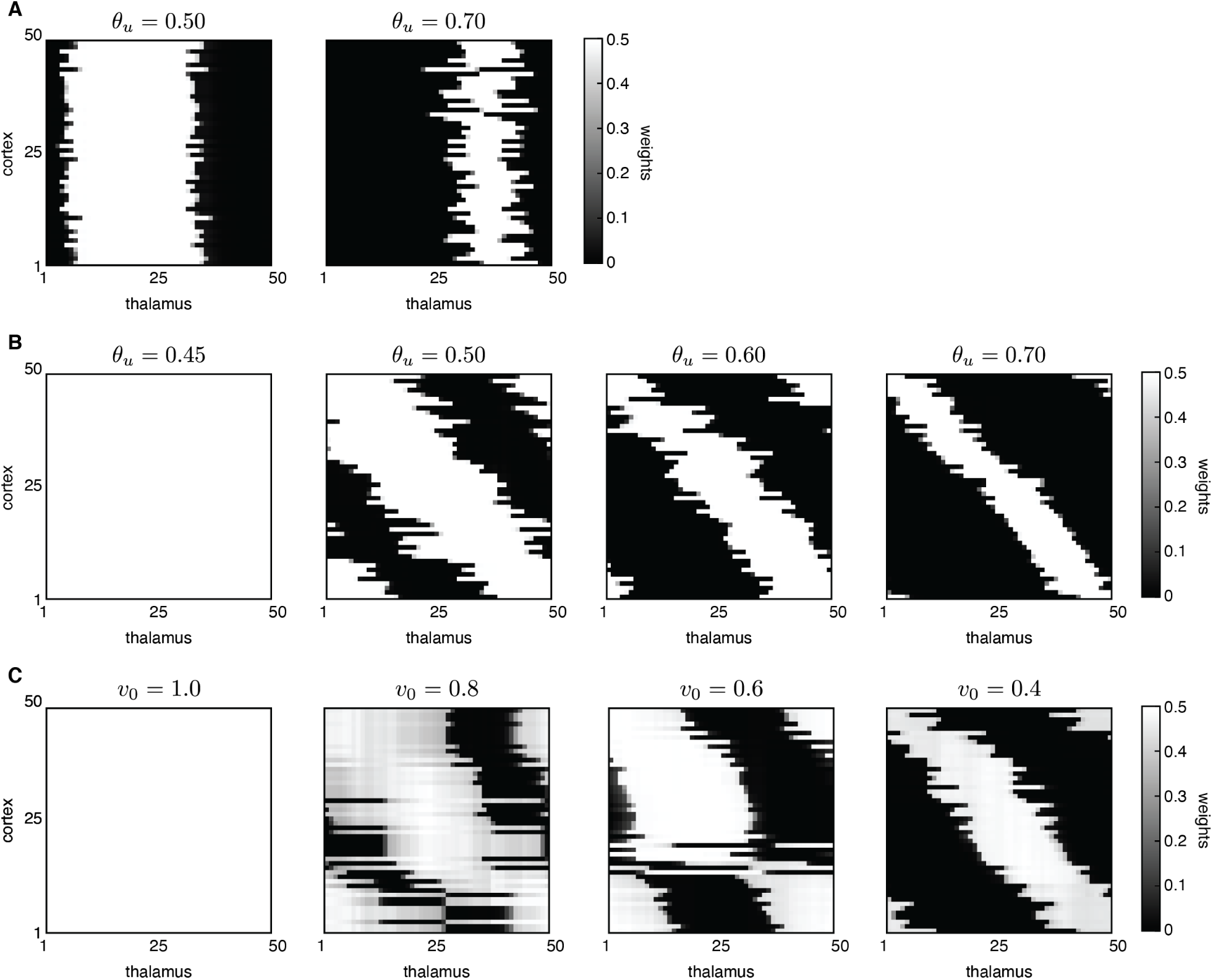
Peripheral L-events generate robust receptive field refinement. **A**. Without the weak bias along the diagonal, selectivity is achieved so that each cortical cell is connected to a few thalamic inputs; however, the same set of input neurons innervates all cortical neurons. Receptive fields were generated with the Hebbian rule, but similar outcomes are obtained with the BCM rule. **B**. Receptive fields generated by the Hebbian rule with varying input threshold *θ*_*u*_. **C**. Receptive fields generated by the BCM rule with varying target output target rate *v*_0_.

**Figure S2.**
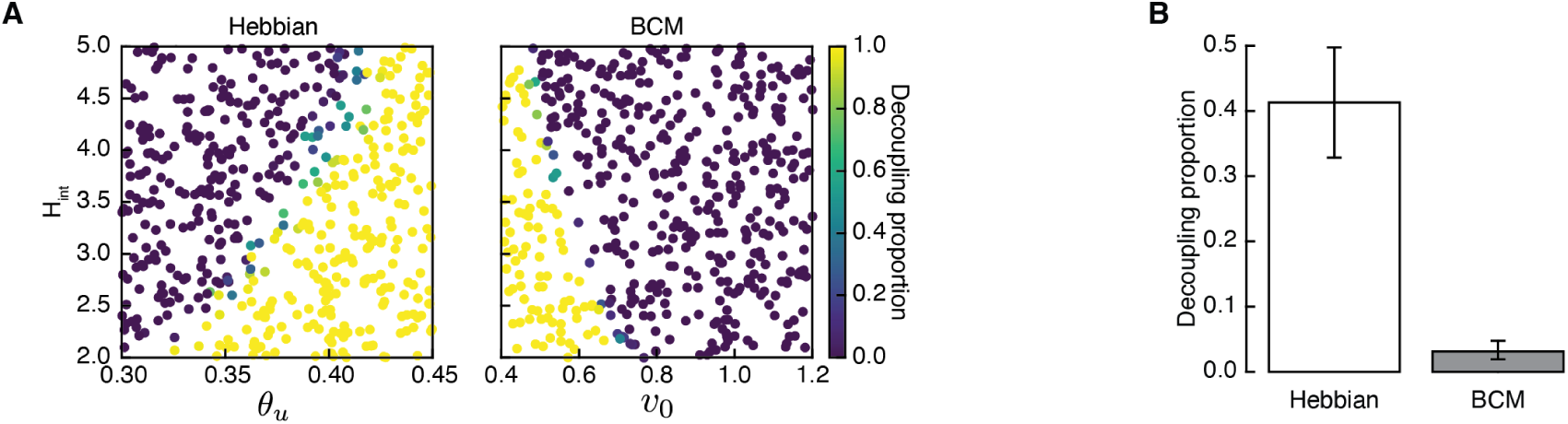
Decoupling of cortical cells is a frequent outcome of H-events in the Hebbian rule. **A**. Proportion of receptive fields that decouple obtained from 500 Monte Carlo simulations for combinations of *H*_int_ and *θ*_*u*_ for the Hebbian rule (left) and *H*_int_ and *v*_0_ for the BCM rule (right). **B**. Average proportion of decoupling of simulation outcomes classified as ‘selective’ in A.

**Figure S3.**
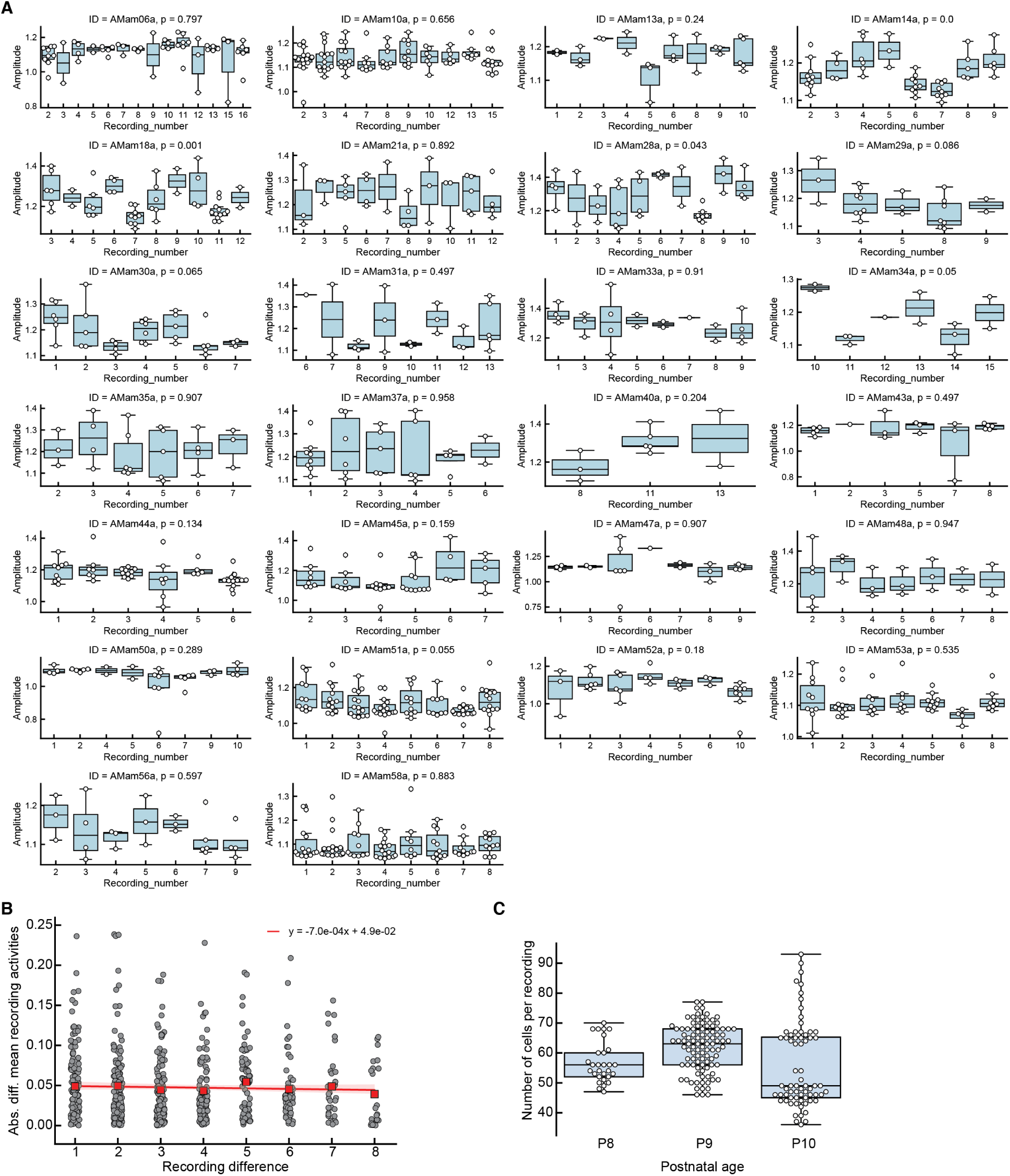
Fluctuations in cortical activity cannot generate correlation. **A**. Spontaneous events have the same mean amplitudes across consecutive recordings (each *∼* 5 mins long) in the same animal (3-14 recordings in 26 animals separated by *<* 5 mins due to experimental constraints on data collection). Each plot corresponds to one animal where each boxplot contains the event amplitudes (both L- and H-events) from each recording. P-values of one-way ANOVA tests of the event amplitudes across recordings are shown for each animal. **B**. Absolute difference in the mean event amplitude as a function of time between recordings. Mean values of each recording difference shown as red squares. The values follow the same distribution (Kruskal-Wallis test p = 0.41). The slope of the linear regression fit (red) is very close to 0, confirming that no clear trend of increasing/decreasing activity exists across recordings. **C**. Number of cells per recording, grouped by different postnatal ages.

**Figure S4.**
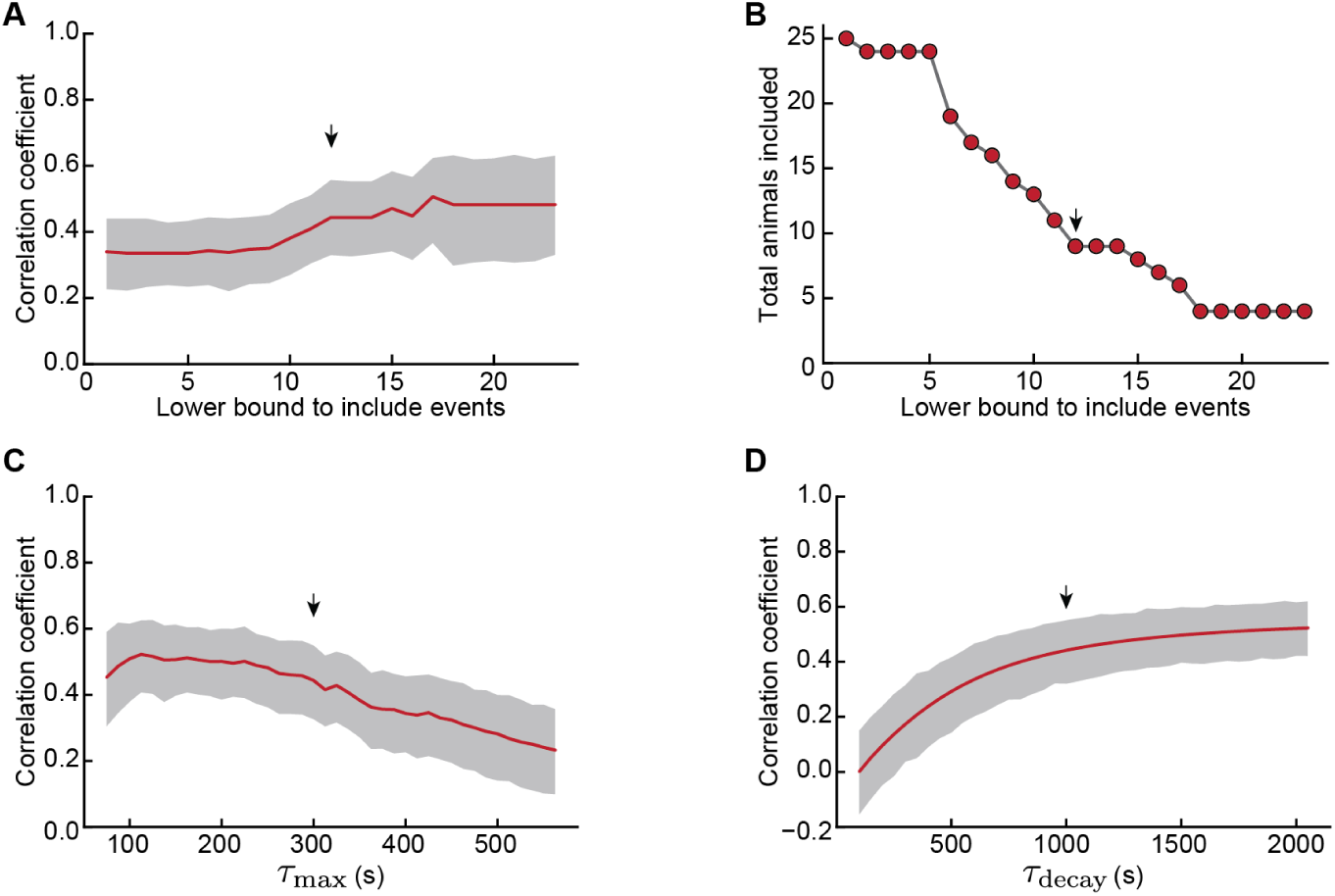
Robustness of correlation under variations in the inclusion criteria, *τ*_max_ and *τ*_decay_. **A**. Correlation coefficient between H-event amplitude and leaky average preceding activity for an increasing lower bound in the number of events necessary to include an animal in the analysis (*τ*_max_ = 300 s and *τ*_decay_ = 1000 s fixed). 95% confidence interval (shaded area) computed through bootstrap regression (Methods). **B**. Corresponding number of animals satisfying the lower bound criteria in A. **C**. Correlation coefficient between H-event amplitude and leaky average preceding activity for varying windows of integration *τ*_max_ (lower bound to include events fixed in 12, *τ*_decay_ = 1000 s). **D**. Correlation coefficient as a function of the exponential decay time constant *τ*_decay_ (lower bound to include events fixed in 12, *τ*_max_ = 300 s). The arrows indicate the parameters of the analysis in Fig. 5, namely a minimal of 12 events per animal, *τ*_max_ = 300 s, *τ*_decay_ = 1000 s.

**Figure S5.**
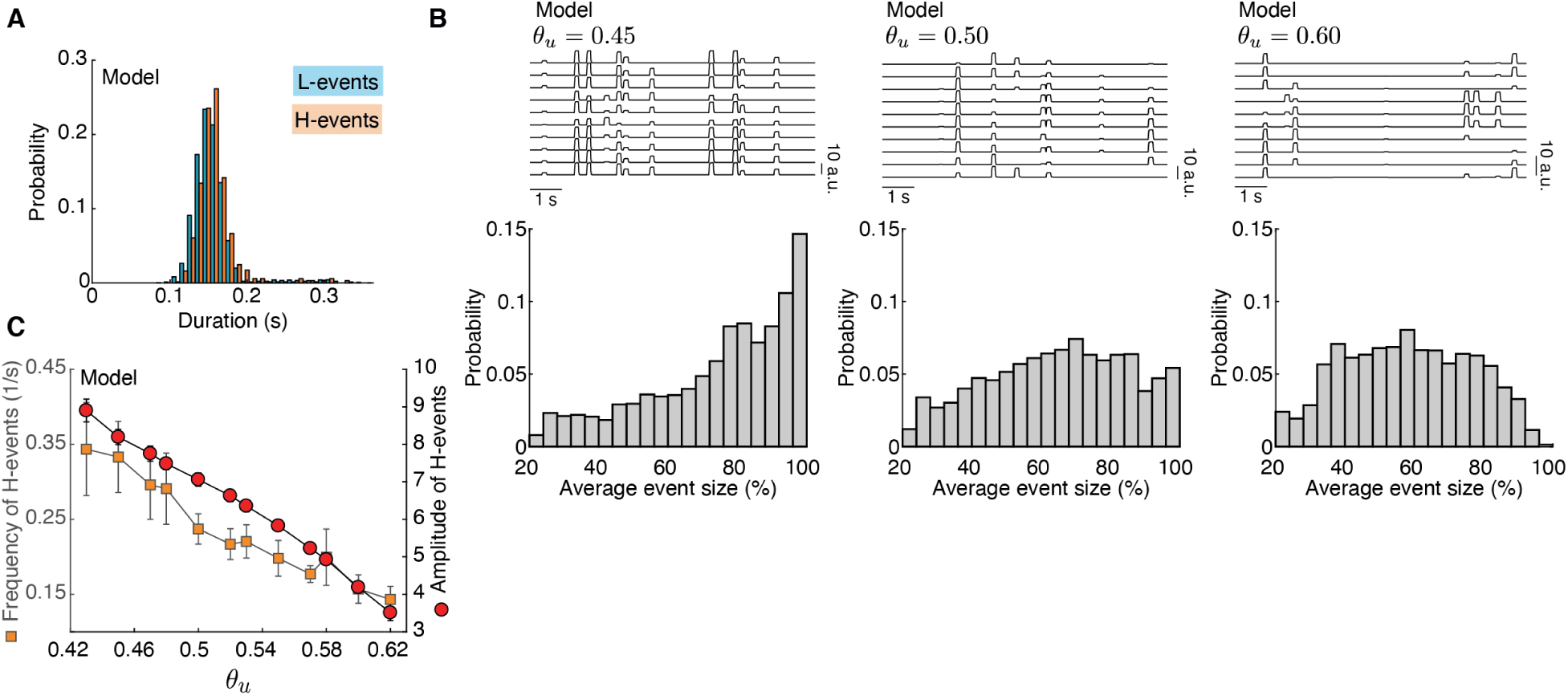
Properties of effective spontaneous events: duration, distribution of event sizes and H-event frequency and amplitude as a function of the input threshold. **A**. The durations of effective L- and H-events in the data are normally distributed with the same parameters. **B**. Top: Sample traces of cortical activity corresponding to Fig. 7D. Bottom: Distribution of spontaneous event sizes for the different input thresholds in the model as Fig. 7D. **C**. Frequencies (squares) and amplitudes (circles) of H-events in the model at different input thresholds (abscissa). H-events are suppressed both in amplitude and frequency at higher input threshold *θ*_*u*_. Bars represent the standard deviation of 10 simulation runs.

